# Novel Automatic Classification of Human Adult Lung Alveolar Type II Cells Infected with SARS-CoV-2 through Deep Transfer Learning Approach

**DOI:** 10.1101/2024.04.22.590420

**Authors:** Turki Turki, Sarah Al Habib, Y-h. Taguchi

## Abstract

SARS-CoV-2 can infect alveoli, inducing a lung injury and thereby impairing the lung function. Healthy alveolar type II (AT2) cells play a major role in lung injury repair as well as keeping alveoli space free from fluids, which is not the case for infected AT2 cells. Unlike previous studies, this novel study aims to automatically differentiate between healthy and infected AT2 cells with SARS-CoV-2 through using efficient AI-based models, which can aid in disease control and treatment. Therefore, we introduce a highly accurate deep transfer learning (DTL) approach that works as follows. First, we downloaded and processed 286 images pertaining to healthy and infected human AT2 (hAT2) cells, obtained from the electron microscopy public image archive. Second, we provided processed images to two DTL computations to induce ten DTL models. The first DTL computation employs five pre-trained models (including DenseNet201 and ResNet152V2) trained on more than million images from ImageNet database to extract features from hAT2 images. Then, flattening and providing the output feature vectors to a trained densely connected classifier with Adam optimizer. The second DTL computation works in a similar manner with a minor difference in which we freeze the first layers for feature extraction in pre-trained models while unfreezing and training the next layers. Compared to TFtDenseNet201, experimental results using five-fold cross-validation demonstrate that TFeDenseNet201 is 12.37 × faster and superior, yielding the highest average ACC of 0.993 (F1 of 0.992 and MCC of 0.986) with statistical significance (*p* < 2.2 × 10^−16^ from a *t*-test).

## 1. Introduction

SARS-CoV-2 is the virus behind causing a respiratory disease, named COVID-19, in which the main target organ is the human lung [1]. When SARS-CoV-2 binds to ACE2 receptor in alveolar type II (AT2) cells of alveoli in lungs, ACE2 receptor becomes occupied, leading to more lung injury and thereby impaired lung function attributed to damaging the alveoli [2-4]. As the number of COVID-19 patients with lung infection significantly outnumbered healthcare workers, researchers developed AI-based techniques to accurately detect lung infection in COVID-19 patients using different imaging modalities. Hussein et al. [5] proposed a custom convolutional neural network (custom-CNN) to accurately classify COVID-19 patients using two previously studied chest X-ray image datasets from Kaggle and GitHub. In the first dataset, the images were categorized into three class labels, including COVID-19, normal (non-COVID-19), and viral pneumonia. For the second dataset, the images were categorized into COVID-19 and normal (non-COVID-19). Their custom-CNN consisted of a series of convolution and max pooling layers, followed by flattening the extracted feature vectors and provided to a densely connected classifier composed of a stack of five dense layers interleaved with three dropout layers and two BatchNormalization layers. Custom-CNN was trained on a random split of the dataset images while testing the performance on the remaining split. Reported results on a random 20% testing split of the first dataset demonstrating that custom-CNN achieved an accuracy of 0.981, precision of 0.976, recall of 0.983, and F1 of 0.973. For the second dataset, the custom-CNN achieved an accuracy of 0.998, a precision of 0.999, a recall of 0.997, and an F1 of 0.998.

To identify pulmonary diseases using X-ray and CT scan images, Abdullahi et al. [6] presented a CNN named PulmoNet, composed of 26 layers using wide residual blocks (of two convolutional layers with dropout layer), followed by a GlobalAveragePooling layer and one dense layer with SoftMax activation to yield predictions. Adam optimizer was used with categorical cross-entropy loss. They formulated the classification problem into a multiclass classification of pulmonary diseases and also reported results for the binary class classification task pertaining to pulmonary diseases as described as follows. The dataset had 16,435 images in which 883 images for bacterial pneumonia, 1,478 images for viral pneumonia, 3,749 for COVID-19, and 10,325 images for healthy cases. They divided the dataset and utilized 85:15 training to testing split ratio. Then, doing a cross-validation based on randomly assigning examples according to the predefined split ratio and averaging results on testing according to five runs of the cross-validation. Particularly, for the task of testing the performance using 4 classes, they achieved an average accuracy of 0.954. For testing 3-class classification pertaining to bacterial pneumonia, COVID-19, and healthy cases, the presented model achieved an average accuracy of 0.954. Also, achieving an average accuracy of 0.994 for the binary class classification pertaining to discrimination between COVID-19 and healthy image cases. For the binary class classification between pneumonia and healthy image cases, their model achieved an average accuracy of 0.983.

Talukder et al. [7] presented a deep transfer learning (DTL) approach to detect COVID-19 using X-ray images working as follows. First, they used two datasets, where the first had 2,000 COVID-19 X-Ray images pertaining to COVID-19 and normal cases and the second had 4,352 Chest X-Ray images pertaining to these four classes: COVID-19, normal, lung opacity, and viral pneumonia. They divided the two datasets and utilized 80:10 training to validation split ratio while the remaining was for testing. Image augmentation was done through operation such as shears, rotation, flipping, and zooming. For DTL, six pre-trained models (Xception, InceptionResNetV2, ResNet50, ResNet50V2, Efficient-NetB0, and EfficientNetB4) were used. Then, freezing first layers in the feature extraction part of pre-trained models and unfreezing next layers, including GlobalAveragePool layer, two BatchNormalization layers, and two dense layers for prediction. Results demonstrated that EfficientNetB4 when applied to 208 testing images from the first dataset achieved the highest accuracy of 1.00. Moreover, EfficientNetB4 when applied to 480 testing images from the second dataset achieved an accuracy of 0.9917 and F1 of 0.9914.

Abdullah et al. [8] presented a hybrid DTL approach to detect COVID-19 using chest X-ray images. First, they downloaded the images from COVID-19 Radiography database at Kaggle, selecting 2413 images pertaining to COVID-19 and 6807 images related to normal cases. They divided the dataset and utilized 70:30 training (including validation) to testing split ratio. To induce the hybrid model, they utilized VGG16 and VGG19 pretrained models in which they applied the feature extraction part of these two pre-trained models to extract features from the image training set. Then, concatenating the output for the two flattened feature vectors, provided as input to train densely connected classifier of two dense layers and one dropout layer. Also, the input feature vectors were provided to train machine learning algorithms including random forest, naive Bayes, KNN, neural network (NN), support vector machines (SVM) with several kernels including linear, sigmoid, and radial. Experimental results applied to the 30% testing data (i.e.,2766 images out of 9,220) demonstrate the superiority of the hybrid DTL approach when coupled with NN achieving the highest MCC of 0.814, followed by SVM with linear kernel that yielded an MCC of 0.805 when compared to existing pre-trained models of densely connected classifier in the binary classification task. Others have proposed deep learning approaches to detect COVID-19 using X-ray and CT images [9-14].

Although these recent studies aimed to detect lung infected with SARS-COV-2 in COVID-19 patients, the novelty in our study is attributed to the summarized contributions as follows:

1. To the best of our knowledge, this is the first time to study lungs infected with SARS-COV-2 at the cellular level within alveoli in human lung using images generated via transmission electron microscopy. Specifically, we downloaded and processed 286 images pertaining to infected and healthy (control) human alveolar type II (hAT2) cells in alveoli from the electron microscopy public image archive (EMPIAR) at https://www.ebi.ac.uk/empiar/EMPIAR-10533/.
2. We formulated the problem as a binary class classification problem and inducing ten DTL models using two DTL computations [15], where in the first DTL computation we apply five pre-trained models (DenseNet201 [16], NasNetMobile [17], ResNet152V2 [18], VGG19 [19], and Xception [20]) to extract features from hAT2 images, followed by flattening the extracted features into feature vectors provided as input to train a densely connected classifier of three layers including two dense layers and one dropout layer. We refer to induced models via the first DTL computation as TFeDenseNet201, TFeNasNetMobile, TFeResNet152V2, TFeVGG19, and TFeXception. For the second DTL computation, we freeze the first layers in the pre-trained models while unfreezing and training the next layers including a densely connected classifier. Induced models via such DTL computation are referred to as TFtDenseNet201, TFtNasNetMobile, TFtResNet152V2, TFtVGG19, and TFtXception.
3. For fairness of performance comparisons among the ten studied DTL models, we evaluated the performance on the whole dataset of 286 images using five-fold cross-validation in which we provided the same training and testing images in each run to each model. Then, reporting the average performance results of the five runs on testing folds and reported the standard deviation.
4. Our conducted experimental study demonstrated that TFeDenseNet201 achieved the highest average ACC of 0.993, the highest F1 of 0.992 and highest MCC of 0.986 when utilizing five-fold cross-validation. Moreover, these performance results were statistically significant (*p* < 2.2 × 10^−16^, obtained from a *t*-test), demonstrating the generalization ability of TFeDenseNet201. In terms of measuring the training running time, TFeDenseNet201 was 12.37 × faster than its peer TFtDenseNet201, induced via the second DTL method. These results demonstrated the feasibility of studied DTL models and promoting TFeDenseNet20 as an assisting AI tool. We provide details about experimental study including processed datasets in Supplementary Materials.

## 2. Materials and Methods

### 2.1. Data Preprocessing

In Figure 1, we present an illustration for hAT2 images employed in this study, obtained from the electron microscopy public image archive (EMPIAR) at https://www.ebi.ac.uk/empiar/EMPIAR-10533/ (accessed on 13 March 2023) and composed of 577 hAT2 images related to uninfected (control) and infected hAT2 cells [21]. The class label distribution consisted of 326 images belonging to control hAT2 cells while the other 251 images belong to infected hAT2 cells. As transmission electron microscopy imaging was employed to generate images, the 577 images had a 4096 × 4224 pixel resolution (i.e., 17.3015Mpx) and were stored as TIFF image files. In our study, we randomly selected and processed 286 TIFF images out of 577 with the use of Image module in python [22], obtaining 286 JPG images with a 256 × 256 pixel resolution for addressing the classification task at hand. It is worth noting that the class distribution was balanced in which 143 images belong to control hAT2 images while the remaining 143 images belong to infected hAT2 images. We provide preprocessed 286 JPG images in Supplementary Dataset.

**Figure 1.**
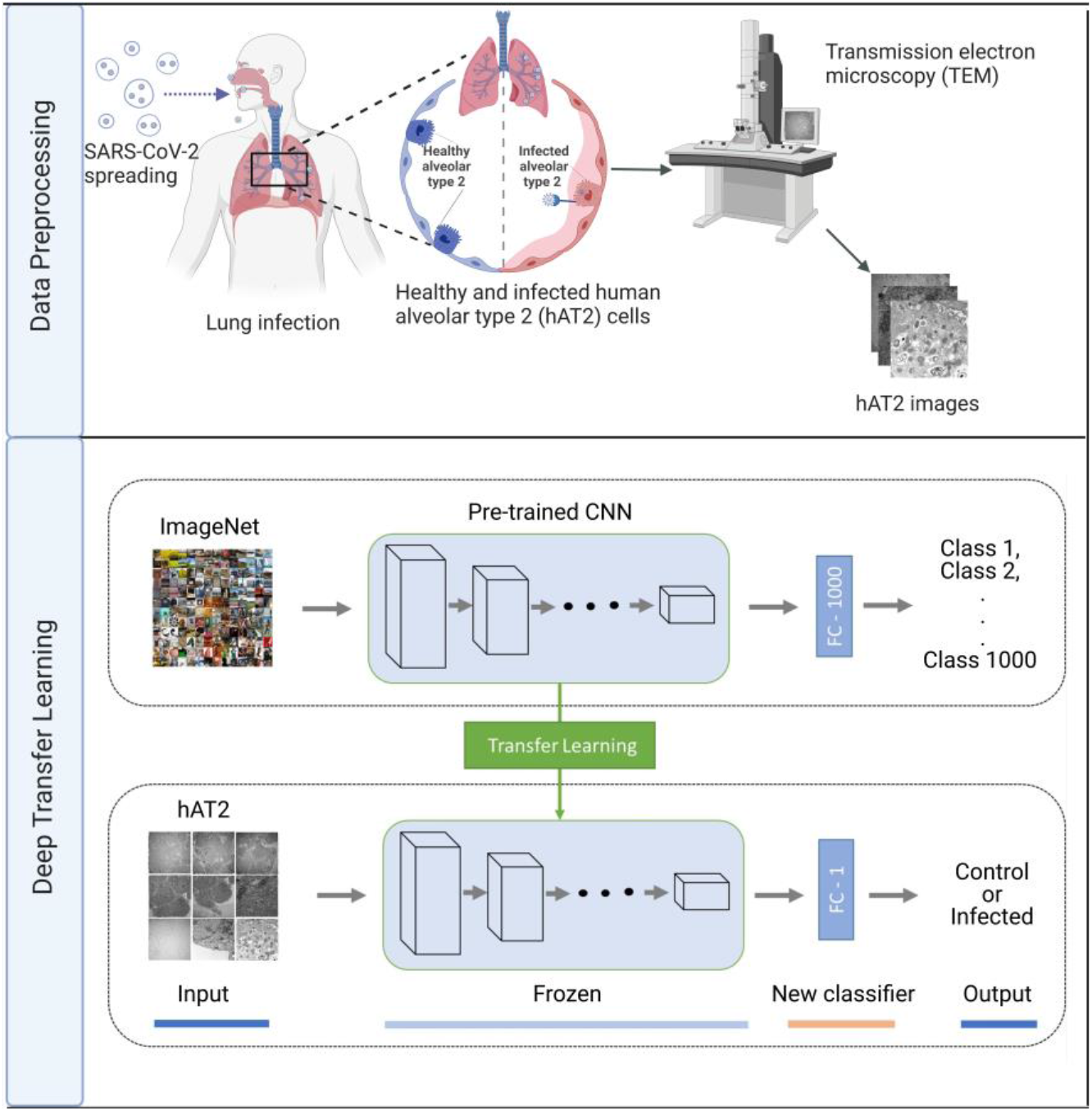
Flowchart showing our deep transfer learning approach for predicting hAT2 cells infected with SARS-CoV-2. Data Preprocessing: To obtain hAT2 images, healthy and infected hAT2 images using transmission electron microscopy (TEM) imaging were downloaded and processed from https://www.ebi.ac.uk/empiar/EMPIAR-10533/. Deep Transfer Learning: hAT2 images are provided to a pre-trained CNN of frozen layers for feature extraction, followed by a trained new classifier to discriminate between healthy and infected hAT2 cells.

### 2.3. Deep Transfer Learning

In Figure 1, we demonstrate how our deep transfer learning (DTL) approach is carried out. Initially, we employed five pre-trained models: DenseNet201, NasNetMobile, ResNet152V2, VGG19, and Xception. As each pre-trained model consists of a feature extraction part (i.e., interleaved convolutional and pooling layers) and densely connected classifier for feature extraction and classification, respectively, we freeze the weights (i.e., keep the weights unaltered) of the features extraction part while modifying the densely connected classifier, dealing with the binary class problem at hand rather than the multiclass classification of 1000 classes. Then, we extract features from hAT2 images via applying the feature extraction part (using unadjusted weights of a pre-trained model), followed by training the densely connected classifiers and performing prediction to unseen hAT2 images. We refer to such DTL models as TFeDenseNet201, TFeNasNetMobile, TFeResNet152V2, TFeVGG19, and TFeXception (see Figure 1). In Table 1, we include information regarding the development of TFe-based models. All models have the same number of unfrozen layers, because we only trained the densely connected classifier (composed of three layers) on extracted features from hAT2 images using pre-trained models. Therefore, the number of non-trainable parameters is 0. As the last 3D tensor output (of shape (8, 8, 2048) for feature extraction was the same for TFeResNet152V2 and TFeXception. Therefore, both had the same number of parameters, as they were flattened into a vector of 131072 elements before feeding into the same densely connected classifier of three layers. We provide details about each TFe-based model in Supplementary TFeModels.

**Table 1.**
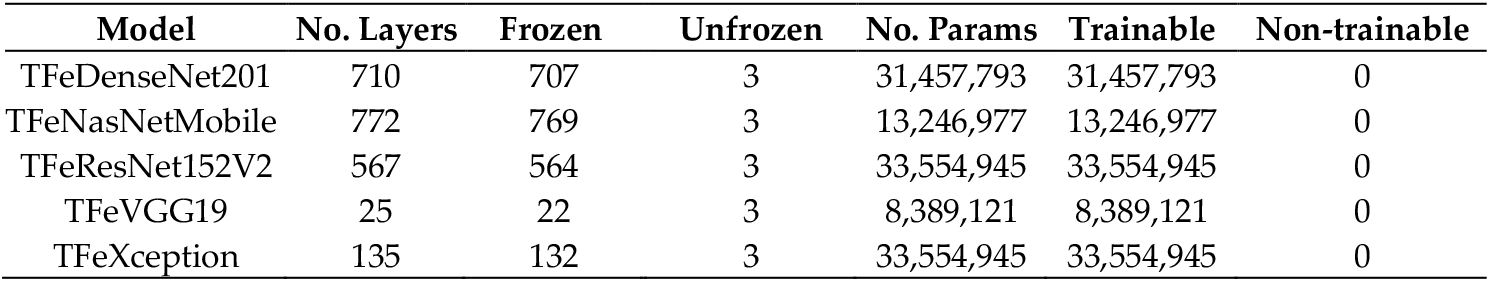
Details about layers and parameters for TFe-based models.

| Model | No. Layers | Frozen | Unfrozen | No. Params | Trainable | Non-trainable |
| --- | --- | --- | --- | --- | --- | --- |
| TFeDenseNet201 | 710 | 707 | 3 | 31,457,793 | 31,457,793 | 0 |
| TFeNasNetMobile | 772 | 769 | 3 | 13,246,977 | 13,246,977 | 0 |
| TFeResNet152V2 | 567 | 564 | 3 | 33,554,945 | 33,554,945 | 0 |
| TFeVGG19 | 25 | 22 | 3 | 8,389,121 | 8,389,121 | 0 |
| TFeXception | 135 | 132 | 3 | 33,554,945 | 33,554,945 | 0 |

For the other DTL models, we freeze the weights of the bottom layers in the feature extraction part while we train the top layers of the feature extraction layers as well as the densely connected classifier. In other words, we unalter weights of the first layers in the feature extraction part while adjusting weights of subsequent layers. Moreover, we modify the densely connected classifier to tackle the binary class classification problem rather than the multiclass classification pertaining to 1000 categories. We refer to models employing such computations as TFtNasNetMobile, TFtResNet152V2, TFtVGG19, and TFtXception (see Figure 2). Table 2 demonstrates the layers and parameters used for TFt-based models. The number of layers including those from pre-trained models in terms of feature extraction plus densely connected classifier of two layers. We report details of each model architecture in Supplementary TFtModels.

**Table 2.** Details about layers and parameters for TFt-based models.

| Model | No. Layers | Frozen | Unfrozen | No. Params | Trainable | Non-trainable |
| --- | --- | --- | --- | --- | --- | --- |
| TFtDenseNet201 | 709 | 649 | 60 | 49,779,777 | 33,605,633 | 16,174,144 |
| TFtNasNetMobile | 771 | 734 | 37 | 17,516,693 | 13,587,361 | 3,929,332 |
| TFtResNet152V2 | 566 | 528 | 38 | 91,886,593 | 48,525,825 | 43,360,768 |
| TFtVGG19 | 24 | 17 | 7 | 28,413,505 | 17,828,35 | 10,585,152 |
| TFtXception | 134 | 129 | 5 | 54,416,425 | 36,718,593 | 17,697,832 |

**Figure 2.**
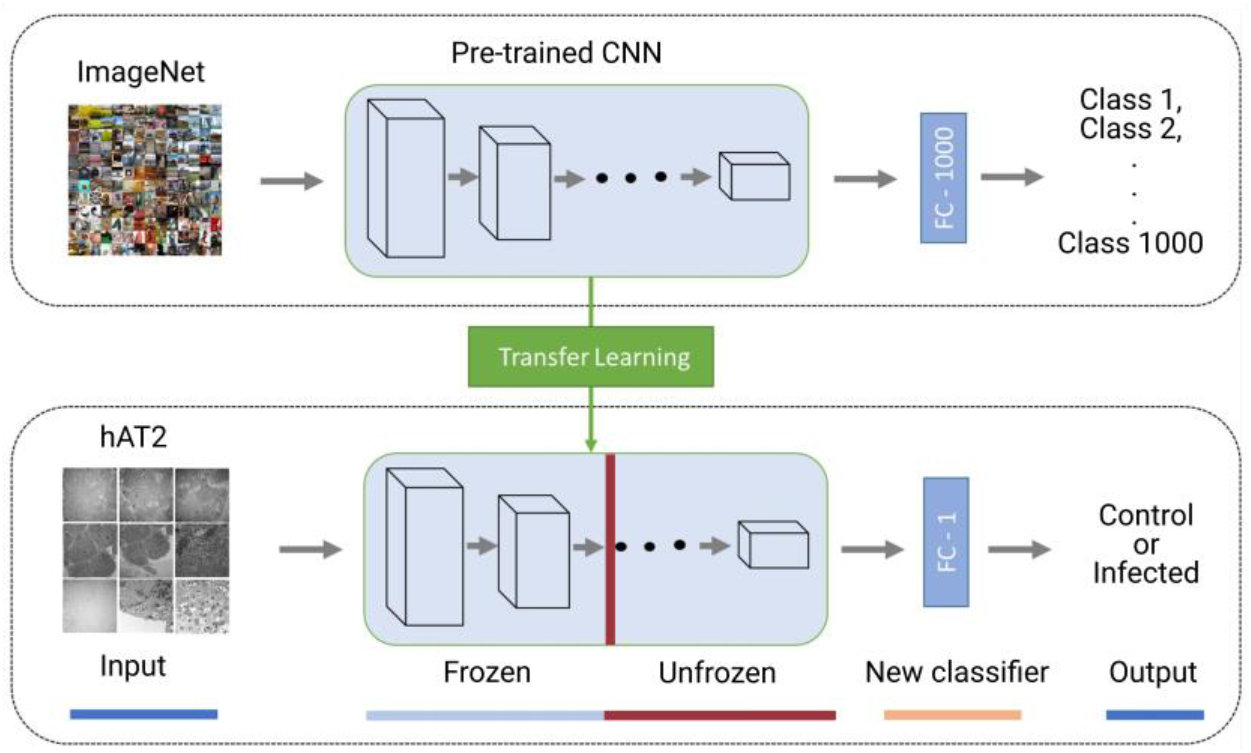
Deep transfer learning composed of a pre-trained CNN with frozen and unfrozen layers for feature extraction, followed by a trained new classifier to discriminate between healthy and infected hAT2 cells.

In the training phase, we employed Adam optimizer with binary crossentropy loss when updating the model parameters. It can be seen that transfer learning is ascribed to the unadjusted weights from pre-trained models that were previously trained on more than million images from ImageNet dataset. In terms of testing the performance to unseen hAT2 images, predictions are mapped to infected if their values are greater than 0.5. Otherwise, predictions are mapped to control.

## 3. Results

### 3.1. Classification Methodology

In this study, we adapted five pre-trained models: DenseNet201, NasNetMobile, ResNet152V2, VGG19, and Xception. Each pre-trained model was trained on over a million images from ImageNet dataset for the multiclass image classification task of 1000 class labels. Then, we developed TFe-based models as follows. We utilized the feature extraction part of pre-trained models by freezing all layers to extract features from hAT2 images, which are then flattened and provided to densely connected classifier (of three layers), trained from scratch for the task of classifying hAT2 cells into control (i.e., uninfected) or infected with SARS-CoV-2. For the TFt-based models, we froze the first layers in the feature extraction part of pre-trained models while unfroze consecutive layers in which the last output was flattened and then provided to densely connected classifier of two layers, trained from scratch to address the binary classification task. For all models, we employed Adam optimizer with binary crossentropy loss function. We assigned the following optimization parameters during the training phase: 0.00001 for the learning rate, 20 for the batch size, and 10 for the number of epochs. We evaluated the performance of each model using accuracy (ACC), F1, and Matthews correlation coefficient (MCC), calculated as follows [23, 24]:

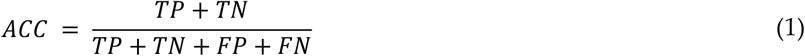

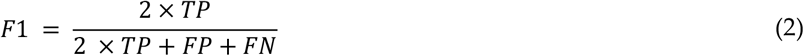

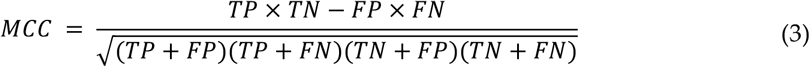

where *TP* is true positive, referred to the number of infected hAT2 images that were correctly predicted as infected. *FN* is false negative, referred to the number of infected hAT2 images that were incorrectly predicted as uninfected (control). *TN* is true negative, referred to the number of control hAT2 images that were correctly predicted as control. *FP* is false positive, referred to the number of control hAT2 images that were incorrectly predicted as infected.

For reporting the performance results on the whole dataset of 286 hAT2 images, we utilized five-fold cross-validation in which we randomly assigned images into five folds. Then, in the first run, we assigned images of the first fold for testing while assigning the remaining examples in the other folds for training, followed by performing prediction to examples in the testing fold and recording results. Such a process was repeated in the remaining four runs in which we recorded the performance on testing folds. Then, taking the average performance calculated during the five runs to be results on five-fold crossvalidation.

### 3.2. Implementation Details

To run the experiments, we used Anaconda distribution, creating an environment with Python (Version 3.10.14) [15], followed by installing the following libraries: cudatoolkit (Version 11.2) [25], cudnn (Version 8.1.0) [26], and tensorflow (Version 2.8) [27]. Then, installing Jupiter notebook [28, 29] to write and execute python code for deep learning models with Keras on our local NVIDIA GeForce RTX 2080Ti GPU with 4352 CUD cores, 11GB of GDDR6 memory, 1545 MHz of boost clock speed, and 14 Gbps of memory clock speed. Other libraries were used for running the experiments, including pandas and NumPy for data processing while matplotlib and Sklearn were utilized for confusion matrix visualization and performance evaluation, respectively [30, 31]. Also, we utilized ggplot2 in R to visualize graphs [32].

### 3.3. Classification Results

#### 3.3.1. Training Results

In Figure 3, we report average training accuracy and loss during running five-fold cross-validation of ten models, trained for 10 epochs. Accuracy is calculated in terms of correctly classified training examples while loss is calculated in terms of the binary crossentropy. These two measures, accuracy, and loss, demonstrate the ability of models to learn from data and thereby will be capable of classifying hAT2 images during testing. At the first epoch, it can be seen that TFeXception generated the highest average accuracy of 0.78 (and lowest average loss of 0.46), followed by TFeDenseNet201, generating an average accuracy (and loss) of 0.68 (and 0.60). TFtDenseNet201 generated the lowest average accuracy (and loss) of 0.42 (and 1.67). At the second epoch, TFeXception adapting more to training data via generating the highest average accuracy of 0.96 (and lowest average loss of 0.13), followed by TFeDenseNet201, achieving and second highest average accuracy of 0.95 (with average loss of 0.17). As the number of epochs increases, the accuracy and loss increase and decrease, respectively, in which all models, except TFeVGG19, achieve an average accuracy above 0.998 and an average loss close to 0. For each epoch, we include average accuracy and loss at Supplementary Epoch.

**Figure 3.**
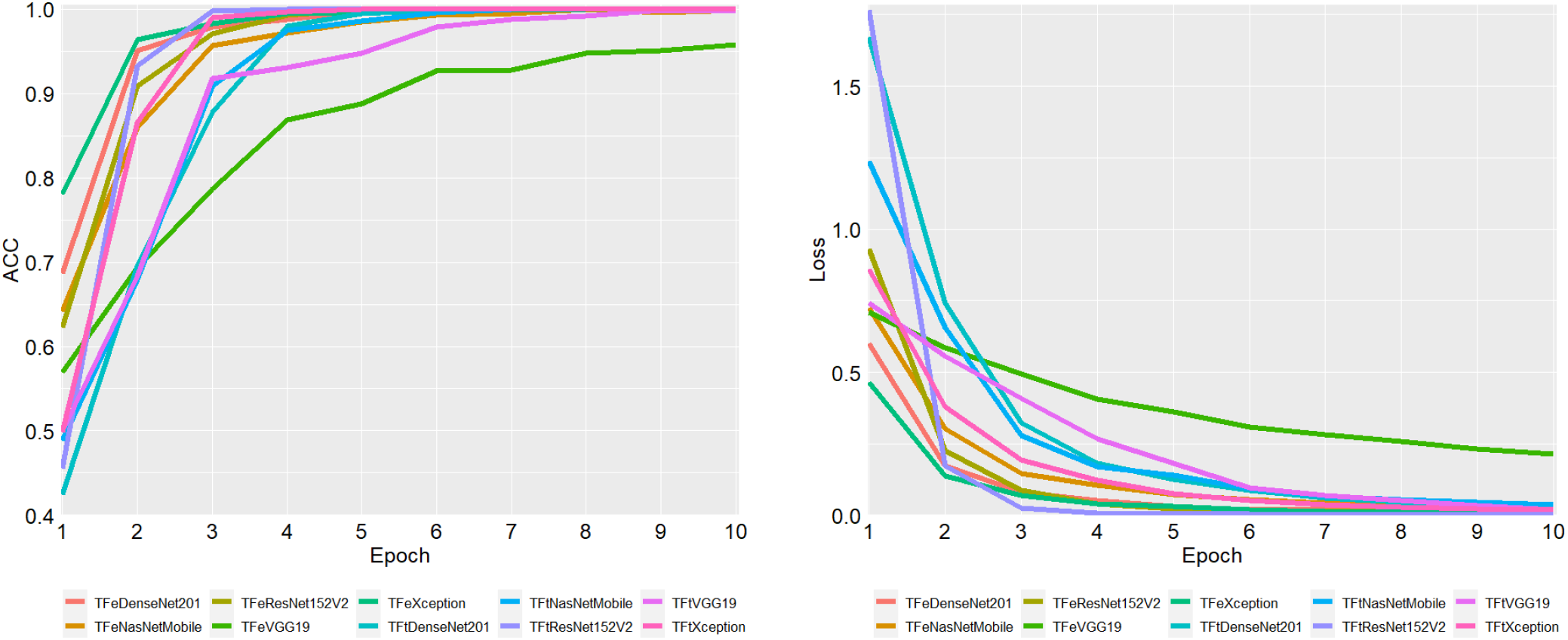
Average ACC and loss of five training folds for each epoch. ACC is accuracy.

Figure 4 illustrates the total running time for each model induction process during running five-fold cross-validation. TFeXception is the fastest. Particularly, TFeXception is 89.03 × faster than its counterpart, TFtXception. The second fastest method is TFeDenseNet201, which is 12.37 × faster than TFtDenseNet201. The third fastest method is TFeResNet152V2, which is 9.26 × faster than TFtResNet152V2. The slowest mode among TFe-based methods is TFeVGG19, which is 11.42 × faster than TFtVGG19. The running time difference between the fastest two methods, TFeXception and TFeDenseNet201, is marginal. Particularly, they differ by less than 1 s. These results demonstrate computational efficiency of TFe-based models, attributed to fewer layers during the training phase.

**Figure 4.**
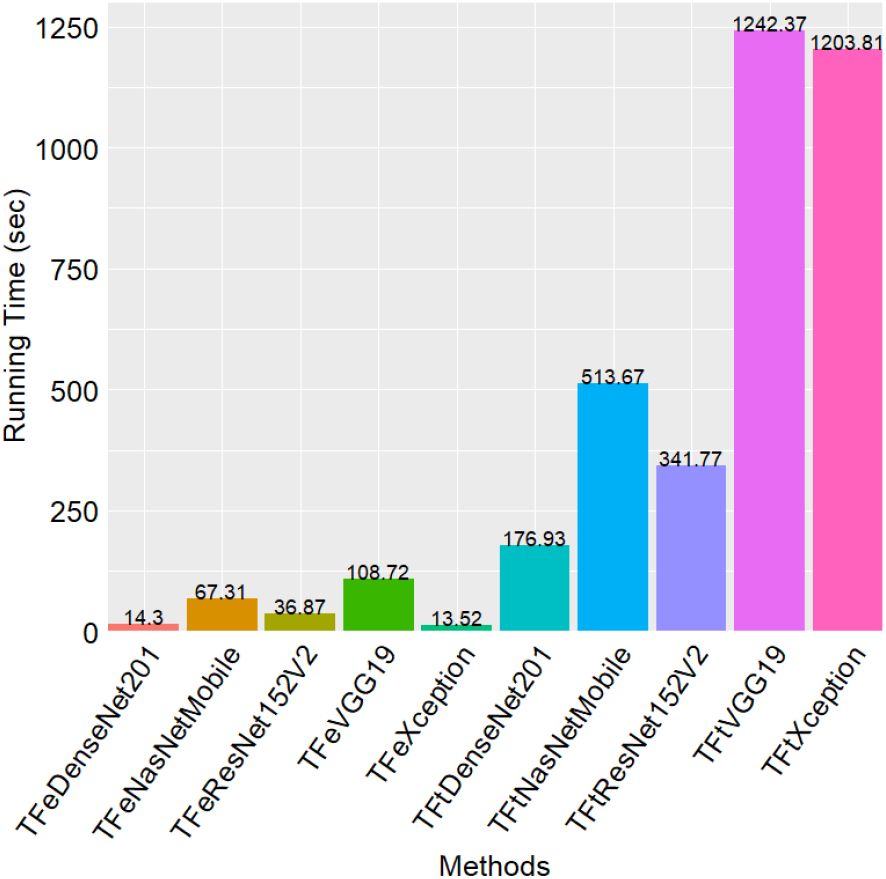
Running time for each method during the training phase in five-fold cross-validation.

#### 3.3.2. Testing Results

In Figure 5, boxplots demonstrate generalization (testing) results of ten models when employing five-fold cross-validation. The median results are displayed by the horizontal bold lines crossing each box. It can be noticed that TFeDenseNet201 is the best performing model, achieving the highest median of 1.00 according to the three performance measures (i.e., ACC, F1, MCC), where the standard deviation σ is 0.009 (0.009, and 0.017) for ACC (F1, and MCC). The second-best performing model is TFeXception, achieving median ACC of 0.982 (σ = 0.009), median F1 of 0.982 (σ = 0.009), and median MCC of 0.966 (σ = 0.017). The third-best performing model is TFtVGG19, achieving median ACC of 0.982 (σ = 0.017), median F1 of 0.982 (σ = 0.018), and median MCC of 0.966 (σ = 0.030). The worst-performing model is TFtResNet152V2, yielding median ACC of 0.785 (σ = 0.064), median F1 of 0.782 (σ = 0.071), median MCC of 0.590 (σ = 0.119). Although TFtVGG19 and TFeXception achieve the same results, the standard deviation for TFtVGG19 demonstrates greater variability in performance results, attributed to degradation in performance in two testing folds when compared to TFeXception. We report median results in Supplementary BoxplotMed.

**Figure 5.**
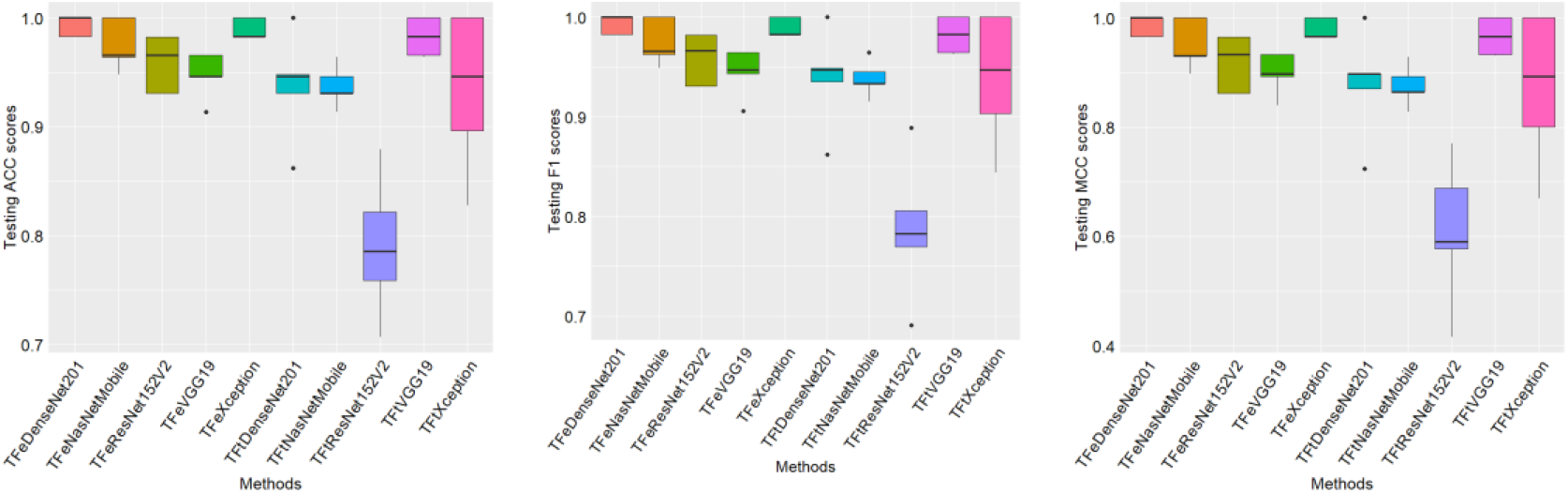
Boxplots show testing (generalization) performance results of ten models during five-fold cross-validation. ACC is accuracy. MCC is Matthews correlation coefficient.

Table 3 records five-fold cross-validation results, which are calculated on the basis of averaging performance results on five testing folds. Also, we record the standard deviation. It can be noticed that TFeDenseNet201 is the best-performing model, achieving the highest average ACC of 0.993 (σ = 0.008), the highest average F1 of 0.992 (σ = 0.009), and the highest average MCC of 0.986 (σ = 0.018). TFeXception is the second-best performing model, yielding an average ACC of 0.989 (σ = 0.008), an average F1 of 0.989 (σ = 0.009), an average MCC of 0.979 (σ = 0.018). TFtVGG19 is the third-best performing model, achieving an average ACC of 0.982 (σ = 0.015), an average F1 of 0.981 (σ = 0.018), an average MCC of 0.966 (σ = 0.033). The worst-performing model is TFtResNet152V2, generating an average ACC of 0.790 (σ = 0.058), an average F1 of 0.787 (σ = 0.071), an average MCC of 0.608 (σ = 0.133). These results demonstrate the superiority of TFe-based models over TFt-based models.

**Table 3.**
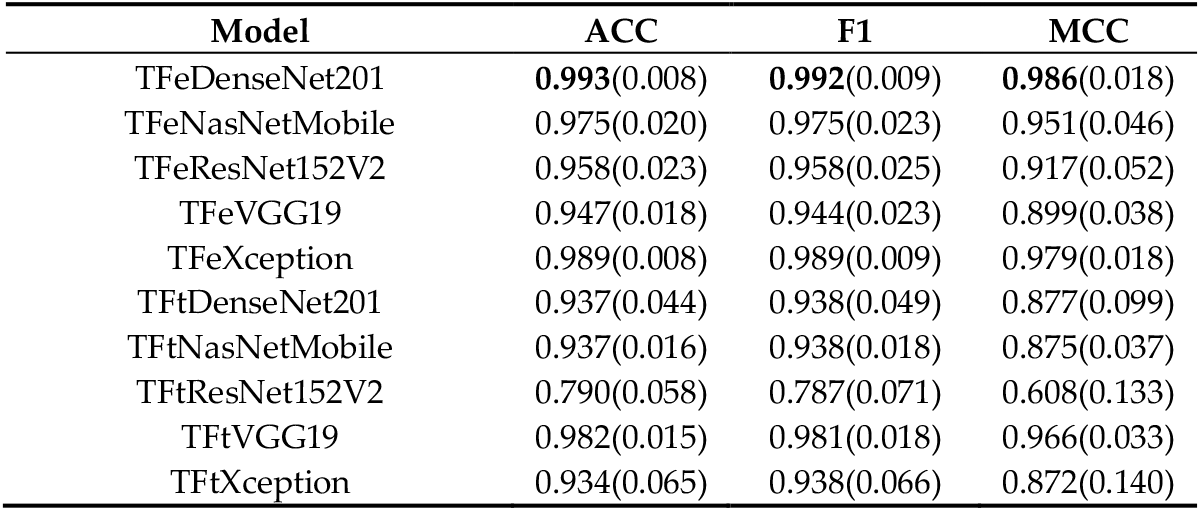
Average performance results during the five-fold cross-validation on test folds for ten models. ACC is accuracy. MCC is Matthews correlation coefficient. Bold refers to a model achieving the highest performance results.

| Model | ACC | F1 | MCC |
| --- | --- | --- | --- |
| TFeDenseNet201 | **0.993(0.008)** | **0.992(0.009)** | **0.986(0.018)** |
| TFeNasNetMobile | 0.975(0.020) | 0.975(0.023) | 0.951(0.046) |
| TFeResNet152V2 | 0.958(0.023) | 0.958(0.025) | 0.917(0.052) |
| TFeVGG19 | 0.947(0.018) | 0.944(0.023) | 0.899(0.038) |
| TFeXception | 0.989(0.008) | 0.989(0.009) | 0.979(0.018) |
| TFtDenseNet201 | 0.937(0.044) | 0.938(0.049) | 0.877(0.099) |
| TFtNasNetMobile | 0.937(0.016) | 0.938(0.018) | 0.875(0.037) |
| TFtResNet152V2 | 0.790(0.058) | 0.787(0.071) | 0.608(0.133) |
| TFtVGG19 | 0.982(0.015) | 0.981(0.018) | 0.966(0.033) |
| TFtXception | 0.934(0.065) | 0.938(0.066) | 0.872(0.140) |

Figure 6 illustrates the combined confusion matrices of test predictions when running five-fold cross-validation. The best-performing model, TFeDeNseNet201, accurately predicted a total of 284 out of 286 hAT2 images while incorrectly predicted 2 images in which the ground truth was infected (positive) and predicted as control (negative), counted as 2 FN. The second-best performing model, TFeXception, accurately predicting 283 hAT2 images while 3 images incorrectly predicted as negative and their actual label is positive, counted as 3 FN. TFtVGG19, the third-best performing model, accurately predicted 281 hAT2 images and incorrectly predicted 5 as negative hAT2 images while their actual label is positive, counted as 5 FN. The worst-performing model, TFtResNet152V2, accurately predicted 226 hAT2 images. Thirty-one hAT2 images were predicted as positive while their label is negative, counted as 31 FP. Moreover, twenty-nine hAT2 images were predicted as negative and their actual label was positive, counted as 29 FN.

**Figure 6.**
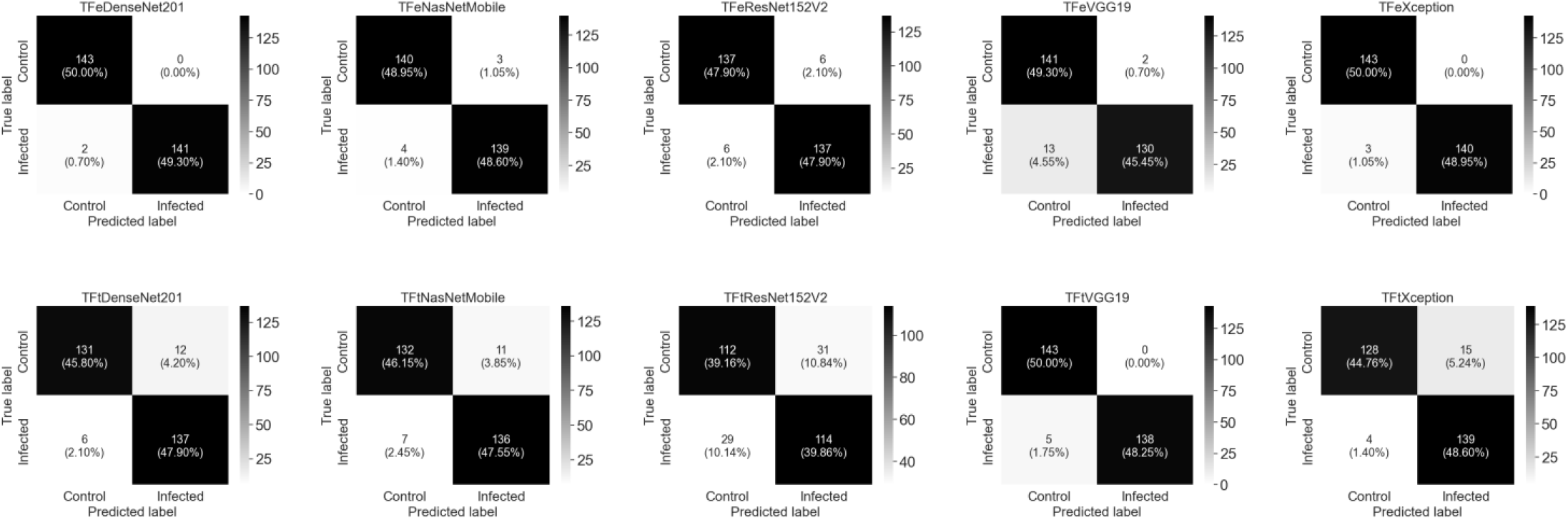
The combined confusion matrices of five testing folds during the running of five-fold cross-validation for ten models.

In Figure 7, we get computational insights for generalization ability of ten studied models, applied to testing. It can be seen from the boxplots and strip charts that prediction differences for TFeDenseNet201 between control hAT2 against infected hAT2 were statistically significant (*P* < 2.2 × 10^−16^, obtained from a *t*-test), suggesting that the TFeDenseNet201 model is a general predictor for control or infected hAT2 prediction. The prediction differences for the second- and third-best performing models (i.e., TFeXception and TFtVGG19) were not significant (*P =* 0.4838 and *P* = 0.1442, respectively, obtained from a *t*-test). These results suggest that these two models are specific, rather than general predictors for hAT2. Other two models, TFeVGG19 and TFtNasNetMobile, exhibited significant prediction differences between control and infected hAT2 (*P =* 3.84 × 10^−3^ and *P* = 4.0 × 10^−2^, respectively, obtained from a *t*- test), although their prediction performance is not outperforming TFeDenseNet201. Prediction differences for all other models between control and infected hAT2 were not significant, suggesting that these models are specific predictors, and we included their *P-*values in Supplementary BoxStripPval.

**Figure 7.**
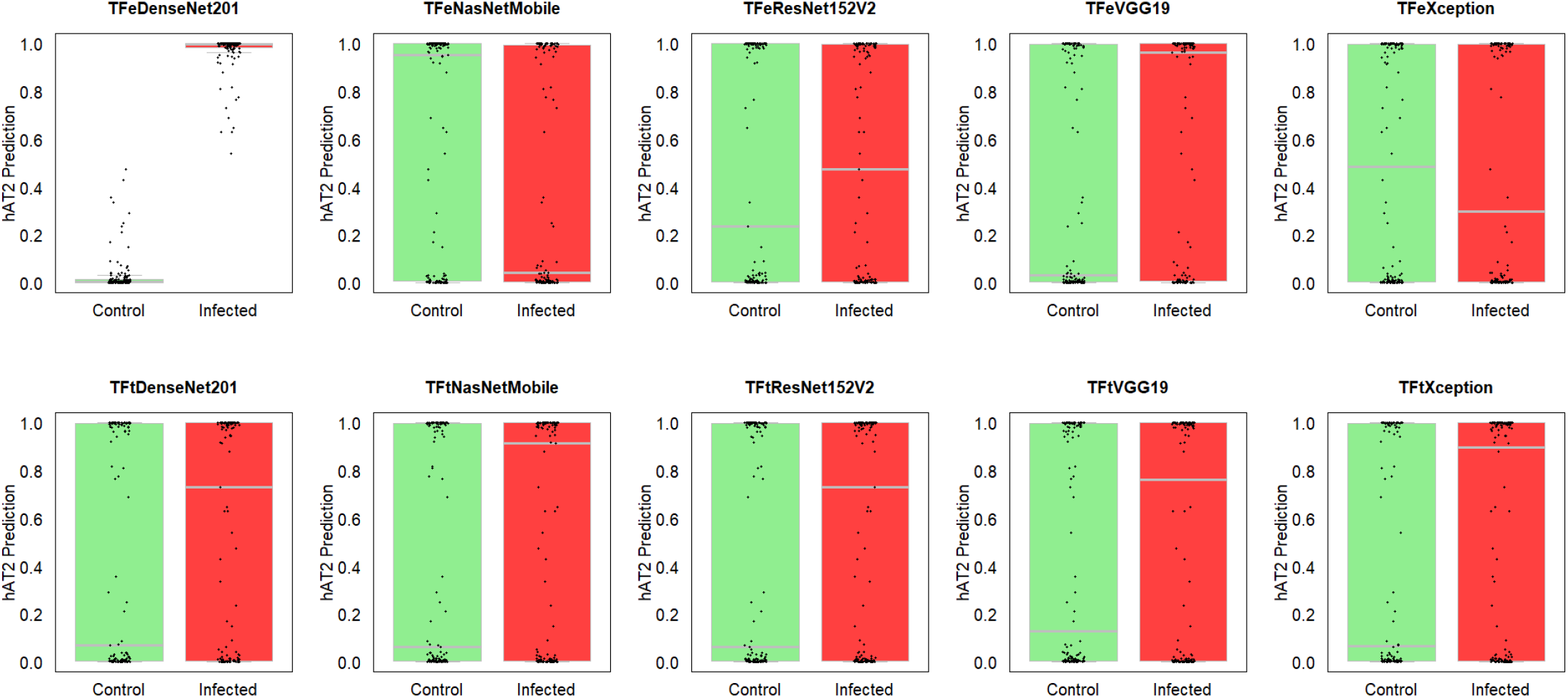
Boxplots and strip charts of predicted hAT2 for control and infected cells on the whole testing folds related to ten models.

## 4. Discussion

To discriminate between control and infected hAT2 cells, our approach consisted of two parts: data preprocessing followed by DTL. In the data preprocessing, we randomly pulled a sample of 286 images pertaining to infected and control hAT2 cells, followed by using Image library in python to convert 286 TIFF images of a 4096 × 4224 pixel resolution to 286 jpg images of a 256 × 256 pixel resolution, which are included in Supplementary Dataset. Then, we conducted two DTL computations using five pre-trained models, namely DenseNet201, NasNetMobile, ResNet152V2, VGG19, and Xception. The first DTL transfers the knowledge by applying the feature extraction part of pre-trained models to hAT2 images in the training set, extracting features that are flattened and provided as input to a densely connected classifier of three layers, trained from scratch to deal with two class labels (i.e., control and infected). We refer to models using this type of transfer learning computations as TFeDenseNet201, TFeNasNetMobile, TFeResNet152V2, TFeVGG19, and TFeXception. The second DTL computation involves the use of the first layers in pre-trained models for feature extraction while training from scratch the next layers including the densely connected classifier, dealing with discriminating between control and infected hAT2 images. Such computation leading to another five DTL models, including TFtDenseNet201, TFtNasNetMobile, TFtResNet152V2, TFtVGG19, and TFtXception.

When we employed five-fold cross-validation, we divided the dataset into training and testing in which the former had examples from four folds while the latter had examples from the remaining one fold. Then, we performed training monitoring the accuracy and loss on the training fold. After finishing from ten epochs, we recorded the loss and accuracy results, and we induced a total of ten DTL models. Then, we applied each DTL to examples in the testing fold and recorded performance results. We repeated such a process for four more runs recording the results, followed by taking the average loss and accuracy (see Figure 3) and reporting average testing results and standard deviation (see Table 3). Experimental results demonstrate the feasibility of DTL in tackling the studied classification task in which TFeDenseNet201 generated the highest average ACC of 0.993, highest average F1 of 0.992, the highest average MCC of 0.986.

It is worth noting that frozen layers in DTL contributed to reducing the number of trainable layers and thereby the mitigation of overfitting. We introduced the dropout layer in the densely connected classifier of TFe-based models to reduce overfitting. In the TFe-based models, we just trained the densely connected classifier composed of three layers while we trained top layers in the feature extraction part in addition to densely connected classifier for TFt-based models. Therefore, we had three unfrozen layers in TFe-based models (see Table 1) while having different number of unfrozen layers for TFt-based models (see Table 2).

In terms of the computational training running time, the two DTL computations were efficient. The DTL computation inducing TFe-based models was more efficient, attributed to the training of just the densely connected classifier. Therefore, it is no surprise that the training time to induce DTL models was longer for computations involving the induction of TFt-based models than computations involved in inducing TFe-based models (see Figure 4). The slowest TFe-based model was TFeVGG19, which was 11.42 × faster than TFtVGG19, the TFt-based model. These demonstrate that DTL computation inducing TFe-based models can efficiently address the task of discriminating between infected and control hAT2 cells.

## 5. Conclusions and Future Work

To address the novel target task of classifying control and infected human alveolar type II (hAT2) cells with SARS-CoV-2, we assess and present ten deep transfer learning (DTL) models derived as follows. First, we downloaded and processed a total of 286 images from the electron microscopy public image archive, pertaining to control and infected hAT2 cells with SARS-CoV-2. Second, we utilized five pre-trained models (DenseNet201, NasNetMobile, ResNet152V2, VGG19, and Xception), previously trained on more than a million images from the ImageNet database. Then, applying the feature extraction part in pre-trained models to extract features from hAT2 images in the training set, followed by performing a flattening step before providing the feature vectors to a modified densely connected classifier with Adam optimizer, trained from scratch to discriminate between control and infected samples. Another DTL computation involves freezing the first layers in the feature extraction part of pre-trained models while unfreezing and training the next layers including a modified densely connected classifier coupled with Adam optimizer. Experimental results on the entire dataset of 286 hAT2 images employing five-fold cross-validation demonstrate (1) the efficiency of TFeDenseNet201 model, which was 12.37 × faster than its counterpart, TFtDenseNet201, during the training time ; (2) the superiority of TFeDenseNet201 achieving the highest average ACC of 0.993 (σ = 0.008), the highest average F1 0.992 (σ = 0.009), and highest average MCC of 0.986 (σ = 0.018), outperforming its counterpart,TFtDenseNet201, that achieved an average ACC of 0.937 (σ = 0.044), an average F1 of 0.938 (σ = 0.049), an average MCC of 0.877(σ = 0.099); (3) significant results achieved via TFeDenseNet201 (*P <* 2.2 × 10^−16^, obtained from a *t*-test) while TFtDenseNet201 did not establish significance (*P* = 0.093, obtained from a *t*-test); (4) the feasibility and reliability of presented TFeDenseNet201 (among other DTL models) as an assisting AI tool for classifying hAT2 cells based on medical imaging.

Future work can include: (1) inducing medical-based imaging models derived from DTL and thereby addressing efficiently different target tasks; (2) developing ensemble models using DTL methods and evaluating their generalization performance; and (3) integrating genomic, clinical information, and features using DTL models to improve the prediction performance in problems from biology and medicine.

## Author Contributions

T.T. conceived and designed the study. T.T. performed the analysis. T.T. and Y-h. T. evaluated the results and discussions. All authors wrote the manuscript. T.T. supervised the study. All authors have read and agreed to the submitted version of the manuscript.

## Funding

This research received no funding.

## Institutional Review Board Statement

Not applicable.

## Informed Consent Statement

Not applicable.

## Data Availability Statement

The hAT2 dataset in this study are available at https://www.ebi.ac.uk/empiar/EMPIAR-10533/, accessed on 13 March 2023.

## Conflicts of Interest

The authors declare no conflicts of interest.

## Abbreviations

The following abbreviations are used in this manuscript:

DL: Deep Learning
DTL: Deep Transfer Learning
DenseNet: Dense Convolutional Network
NasNet: Neural Architecture Search Network
ResNet: Residual Neural Network
VGG Visual: Geometry Group
Xception: Extreme Inception
CNN: Convolutional Neural Network
hAT2: Human Alveolar Type II
SARS-CoV-2: Severe Acute Respiratory Syndrome Coronavirus 2
COVID-19: Coronavirus Disease 19
Adam: Adaptive Moment Estimation
EMPIAR: Electron Microscopy Public Image Archive
CT: Computerized Tomography
TEM: Transmission Electron Microscope
ACC: Accuracy
MCC: Matthews Correlation Coefficient

## Notes

### Competing Interest Statement

The authors have declared no competing interest.

## References

1. Van Slambrouck, J., et al., Visualising SARS-CoV-2 infection of the lung in deceased COVID-19 patients. EBioMedicine, 2023. 92.

2. Gerard, L., et al., Increased angiotensin-converting enzyme 2 and loss of alveolar type II cells in COVID-19–related acute respiratory distress syndrome. American journal of respiratory and critical care medicine, 2021. 204(9): p. 1024–1034.

3. Taguchi, Y. and T. Turki, A new advanced in silico drug discovery method for novel coronavirus (SARS-CoV-2) with tensor decomposition-based unsupervised feature extraction. PloS one, 2020. 15(9): p. e0238907.

4. Kathiriya, J.J., et al., Human alveolar type 2 epithelium transdifferentiates into metaplastic KRT5+ basal cells. Nature cell biology, 2022. 24(1): p. 10–23.

5. Hussein, A.M., et al., Auto-detection of the coronavirus disease by using deep convolutional neural networks and X-ray photographs. Scientific reports, 2024. 14(1): p. 534.

6. Abdulahi, A.T., et al., PulmoNet: a novel deep learning based pulmonary diseases detection model. BMC Medical Imaging, 2024. 24(1): p. 51.

7. Talukder, M.A., et al., Empowering covid-19 detection: Optimizing performance through fine-tuned efficientnet deep learning architecture. Computers in Biology and Medicine, 2024. 168: p. 107789.

8. Abdullah, M., B. Kedir, and T.T. Takore, A Hybrid Deep Learning CNN model for COVID-19 detection from chest X-rays. Heliyon, 2024.

9. Haennah, J.J., C.S. Christopher, and G.G. King, Prediction of the COVID disease using lung CT images by deep learning algorithm: DETS-optimized Resnet 101 classifier. Frontiers in Medicine, 2023. 10: p. 1157000.

10. Celik, G., Detection of Covid-19 and other pneumonia cases from CT and X-ray chest images using deep learning based on feature reuse residual block and depthwise dilated convolutions neural network. Applied Soft Computing, 2023. 133: p. 109906.

11. Park, D., et al., Development and validation of a hybrid deep learning–machine learning approach for severity assessment of COVID-19 and other pneumonias. Scientific Reports, 2023. 13(1): p. 13420.

12. Okada, N., et al., “KAIZEN” method realizing implementation of deep-learning models for COVID-19 CT diagnosis in real world hospitals. Scientific Reports, 2024. 14(1): p. 1672.

13. Salama, G.M., A. Mohamed, and M.K. Abd-Ellah, COVID-19 classification based on a deep learning and machine learning fusion technique using chest CT images. Neural Computing and Applications, 2024. 36(10): p. 5347–5365.

14. Ju, H., et al., CODE-NET: A deep learning model for COVID-19 detection. Computers in Biology and Medicine, 2024: p. 108229.

15. Chollet, F., Deep learning with Python. 2021: Simon and Schuster.

16. Huang, G., et al. Densely connected convolutional networks. in Proceedings of the IEEE conference on computer vision and pattern recognition. 2017.

17. Zoph, B., et al. Learning transferable architectures for scalable image recognition. in Proceedings of the IEEE conference on computer vision and pattern recognition. 2018.

18. He, K., et al. Identity mappings in deep residual networks. in Computer Vision–ECCV 2016: 14th European Conference, Amsterdam, The Netherlands, October 11–14, 2016, Proceedings, Part IV 14. 2016. Springer.

19. Simonyan, K. and A. Zisserman, Very Deep Convolutional Networks for Large-Scale Image Recognition, in 3rd International Conference on Learning Representations. 2015: San Diego, CA, USA.

20. Chollet, F. Xception: Deep learning with depthwise separable convolutions. in Proceedings of the IEEE conference on computer vision and pattern recognition. 2017.

21. Youk, J., et al., Three-dimensional human alveolar stem cell culture models reveal infection response to SARS-CoV-2. Cell Stem Cell, 2020. 27(6): p. 905-919. e10.

22. Clark, A., Pillow (pil fork) documentation. readthedocs, 2015.

23. Alghamdi, S. and T. Turki, A novel interpretable deep transfer learning combining diverse learnable parameters for improved T2D prediction based on single-cell gene regulatory networks. Scientific Reports, 2024. 14(1): p. 4491.

24. Turki, T. and Z. Wei, Boosting support vector machines for cancer discrimination tasks. Computers in biology and medicine, 2018. 101: p. 236–249.

25. Fatica, M. CUDA toolkit and libraries. in 2008 IEEE hot chips 20 symposium (HCS). 2008. IEEE.

26. Chetlur, S., et al., cudnn: Efficient primitives for deep learning. arXiv preprint 1410.0759, 2014.

27. Abadi, M., et al. TensorFlow: a system for Large-Scale machine learning. in 12th USENIX symposium on operating systems design and implementation (OSDI 16). 2016.

28. Granger, B.E. and F. Pérez, Jupyter: Thinking and storytelling with code and data. Computing in Science & Engineering, 2021. 23(2): p. 7–14.

29. Perez, F. and B.E. Granger, Project Jupyter: Computational narratives as the engine of collaborative data science. Retrieved September, 2015. 11(207): p. 108.

30. McKinney, W., Python for data analysis. 2022: “O’Reilly Media, Inc.”.

31. Pedregosa, F., et al., Scikit-learn: Machine learning in Python. the Journal of machine Learning research, 2011. 12: p. 2825–2830.

32. Wickham, H., W. Chang, and M.H. Wickham, Package ‘ggplot2’. Create elegant data visualisations using the grammar of graphics. Version, 2016. 2(1): p. 1–189.

