## Supplementary figures and images for "Novel Automatic Classification of Human Adult Lung Alveolar Type II Cells Infected with SARS-CoV-2 through Deep Transfer Learning Approach"

### c_7.jpg

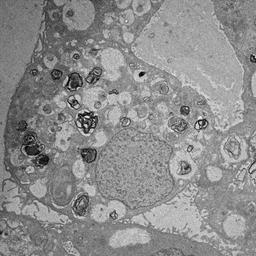

### c_8.jpg

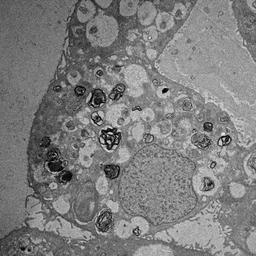

### c_9.jpg

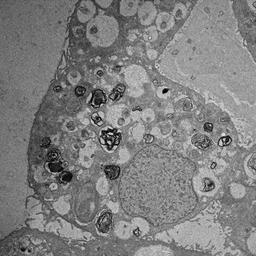

### c_60.jpg

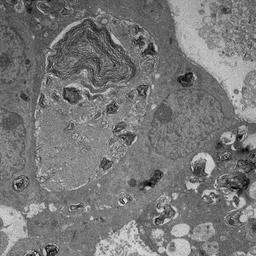

### c_61.jpg

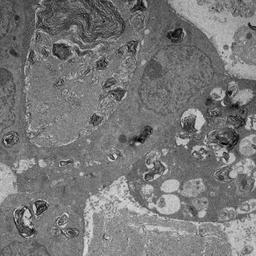

### c_62.jpg

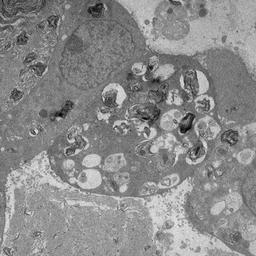

### c_63.jpg

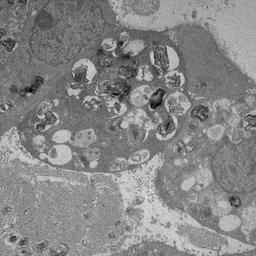

### c_64.jpg

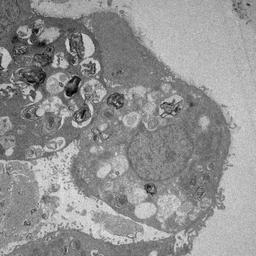

### c_65.jpg

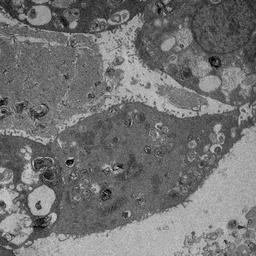

### c_66.jpg

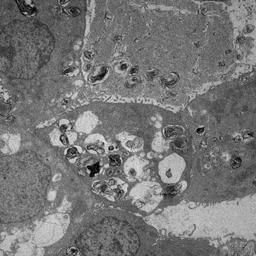

### c_67.jpg

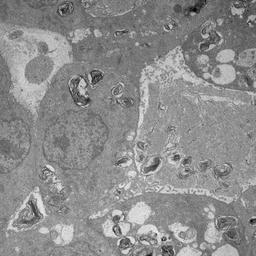

### c_68.jpg

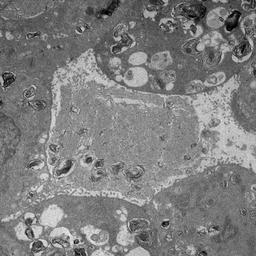

### c_69.jpg

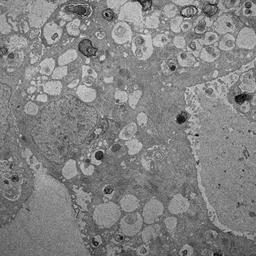

### c_70.jpg

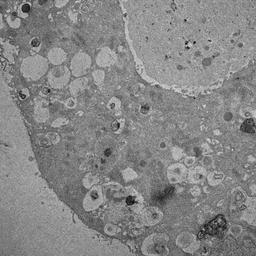

### c_71.jpg

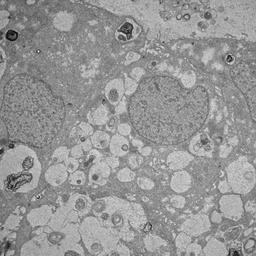

### c_72.jpg

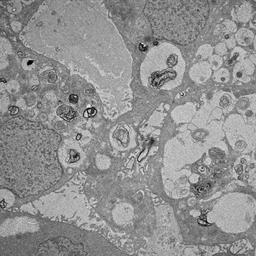

### c_73.jpg

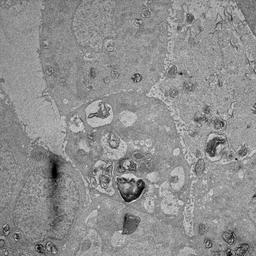

### c_74.jpg

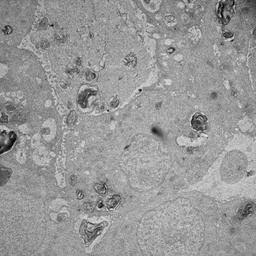

### c_75.jpg

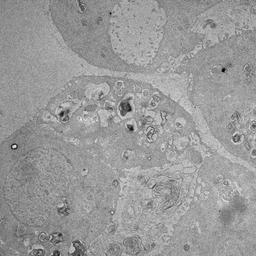

### c_76.jpg

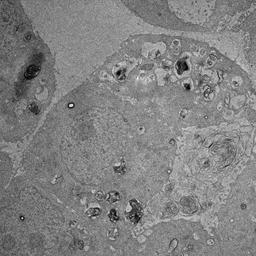

### c_77.jpg

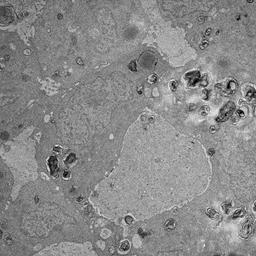

### c_78.jpg

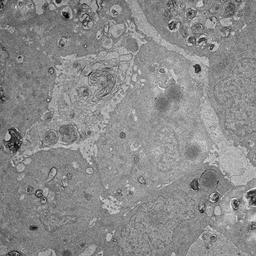

### c_79.jpg

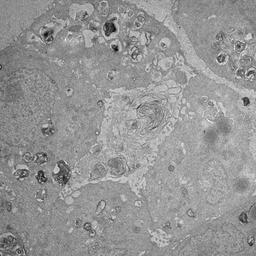

### c_80.jpg

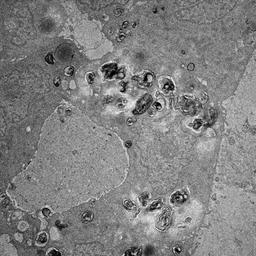

### c_81.jpg

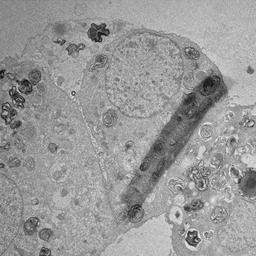

### c_82.jpg

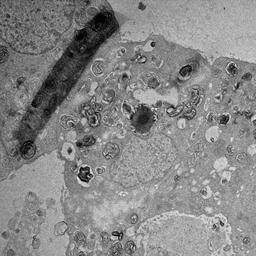

### c_83.jpg

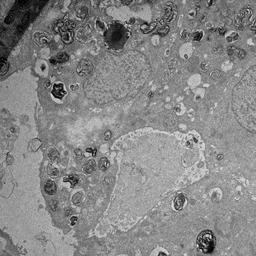

### c_84.jpg

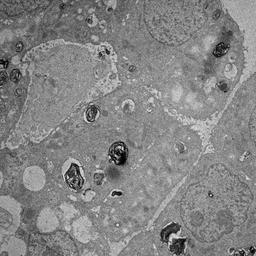

### c_85.jpg

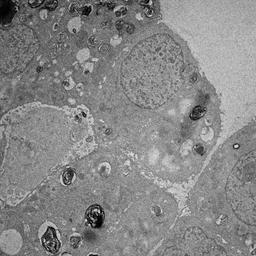

### c_86.jpg

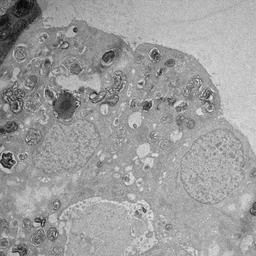

### c_87.jpg

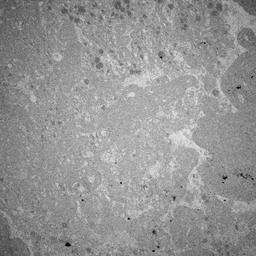

### c_88.jpg

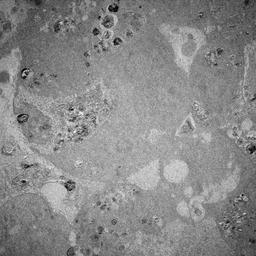

### c_89.jpg

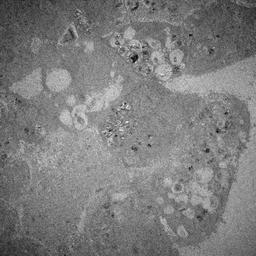

### c_90.jpg

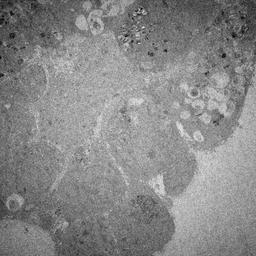

### c_91.jpg

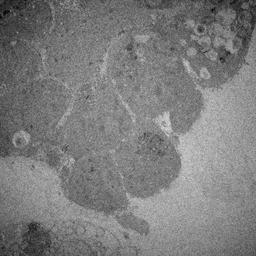

### c_92.jpg

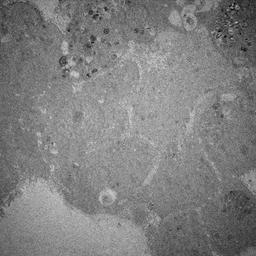

### c_93.jpg

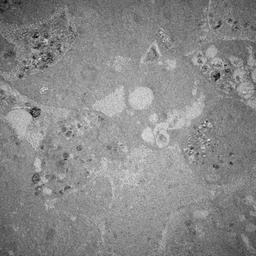

### c_94.jpg

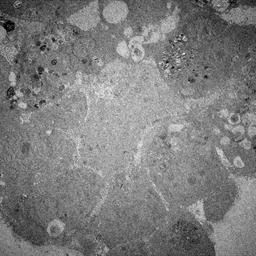

### c_95.jpg

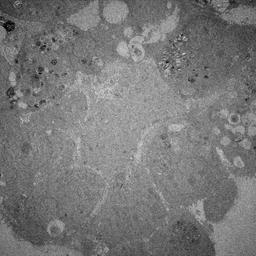

### c_96.jpg

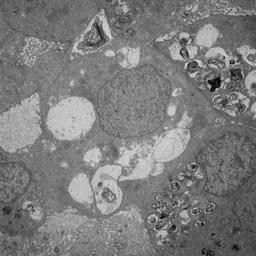

### c_97.jpg

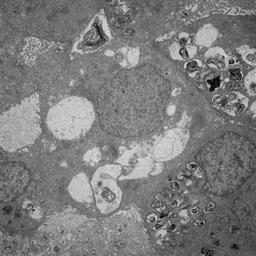

### c_98.jpg

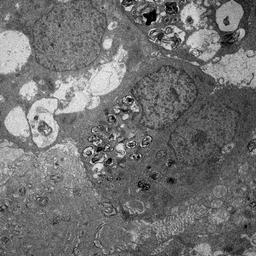

### c_99.jpg

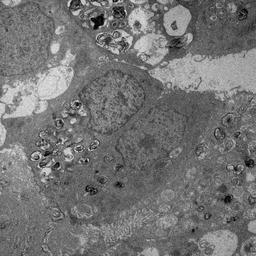

### i_1.jpg

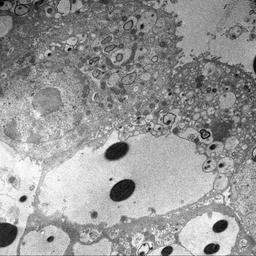

### i_10.jpg

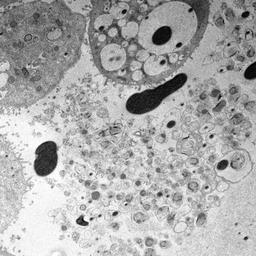

### i_11.jpg

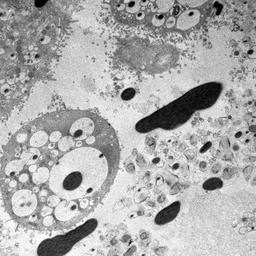

### i_12.jpg

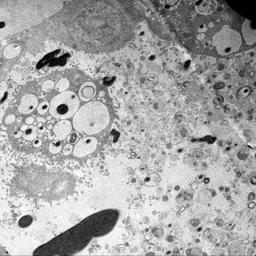

### i_13.jpg

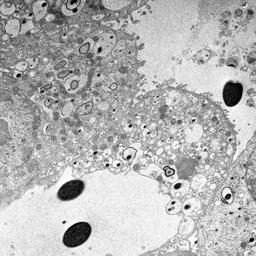

### i_14.jpg

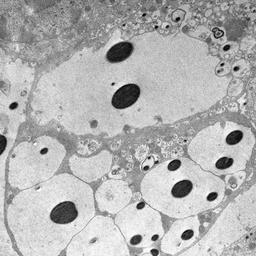

### i_15.jpg

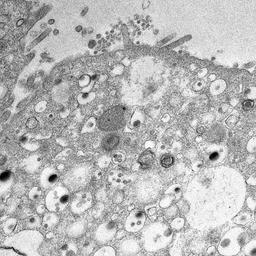

### i_16.jpg

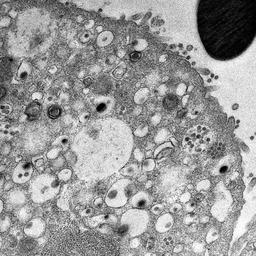

### i_17.jpg

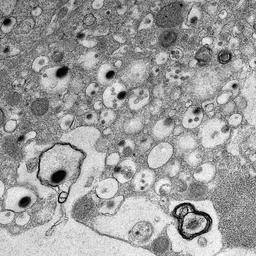

### i_18.jpg

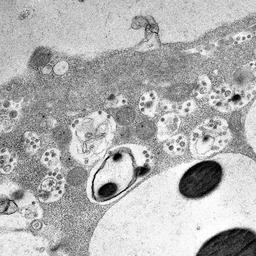

### i_19.jpg

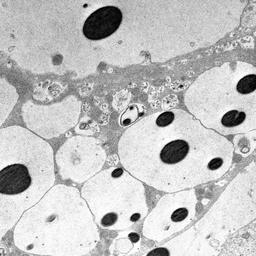

### i_100.jpg

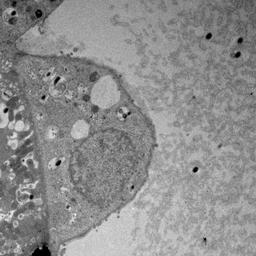

### i_101.jpg

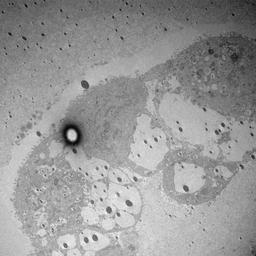

### i_102.jpg

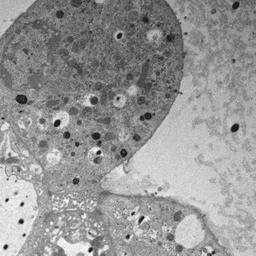

### i_103.jpg

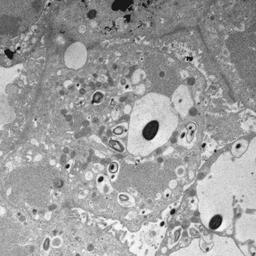

### i_104.jpg

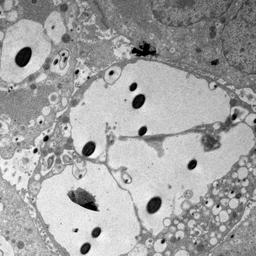

### i_105.jpg

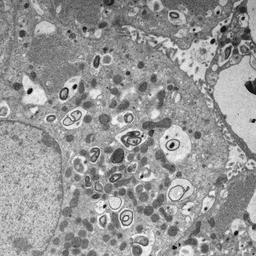

### i_106.jpg

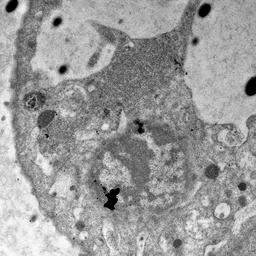

### i_107.jpg

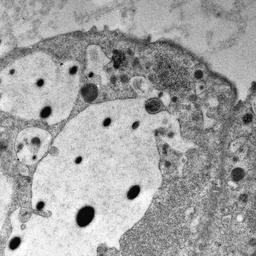

### i_108.jpg

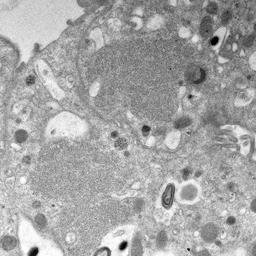

### i_109.jpg

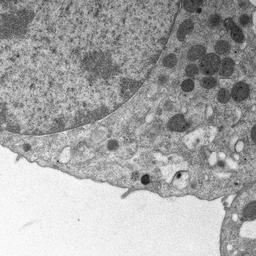

### i_110.jpg

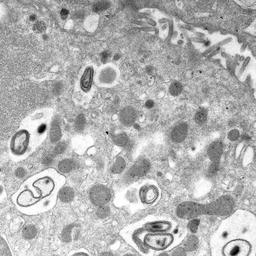

### i_111.jpg

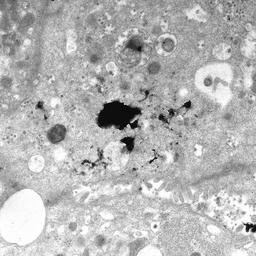

### i_112.jpg

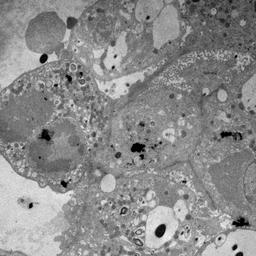

### i_113.jpg

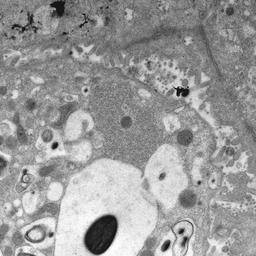

### i_114.jpg

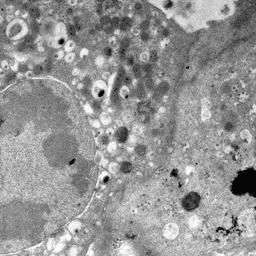

### i_115.jpg

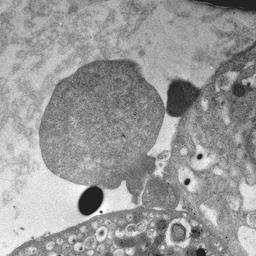

### i_116.jpg

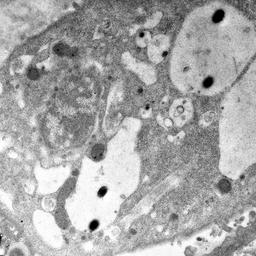

### i_117.jpg

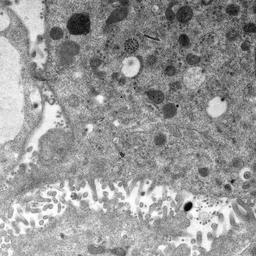

### i_118.jpg

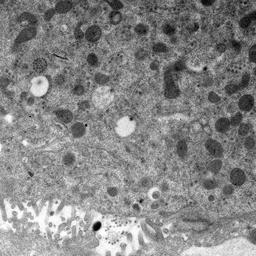

### i_119.jpg

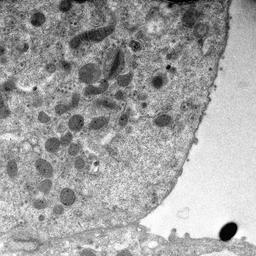

### i_120.jpg

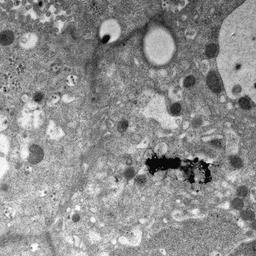

### i_121.jpg

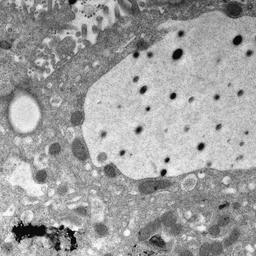

### i_122.jpg

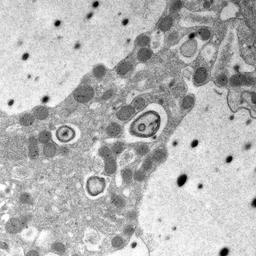

### i_123.jpg

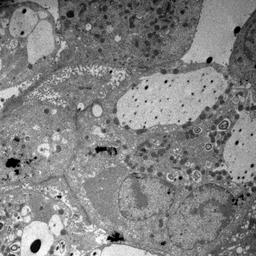

### i_124.jpg

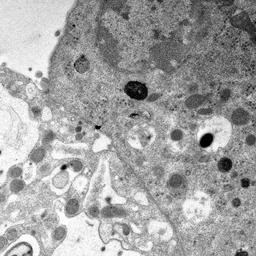

### i_125.jpg

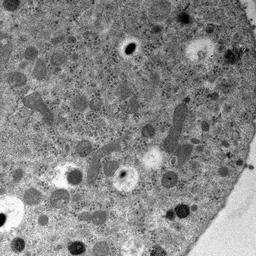

### i_126.jpg

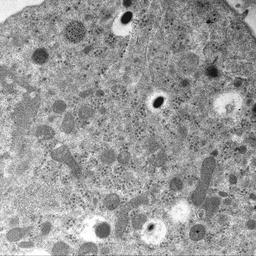

### i_127.jpg

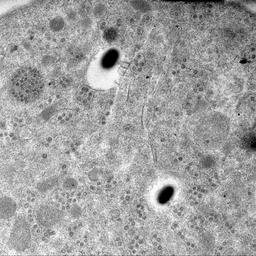

### i_128.jpg

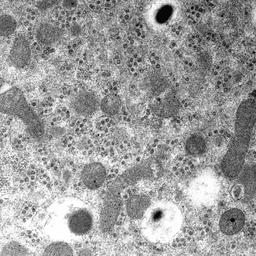

### i_129.jpg

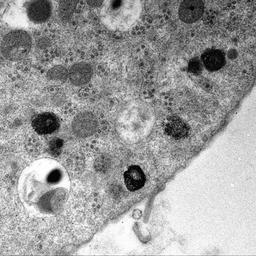

### i_130.jpg

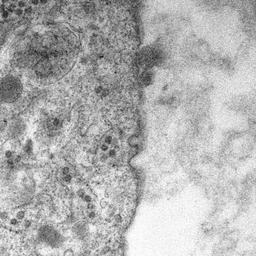

### i_131.jpg

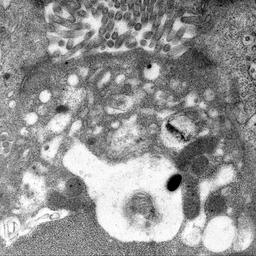

### i_132.jpg

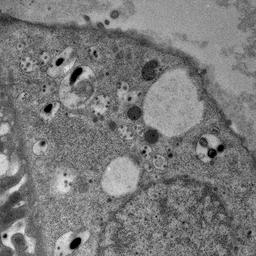

### i_133.jpg

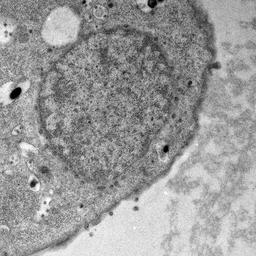

### i_134.jpg

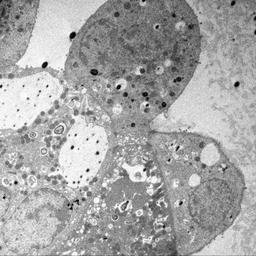

### i_135.jpg

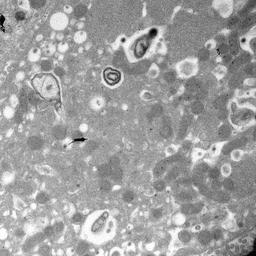

### i_136.jpg

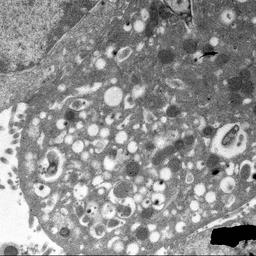

### i_137.jpg

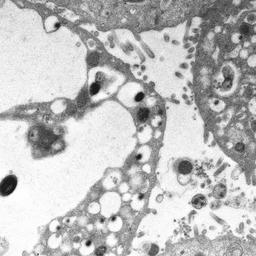

### i_138.jpg

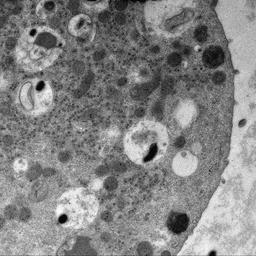

### i_139.jpg

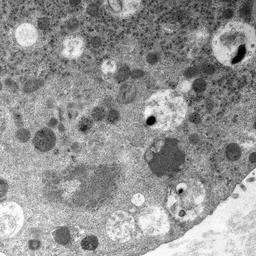

### i_140.jpg

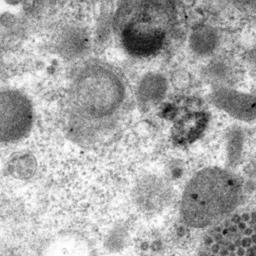

### i_141.jpg

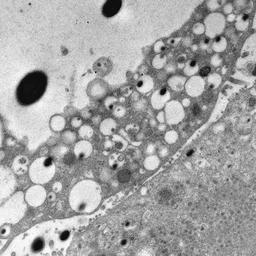

### i_142.jpg

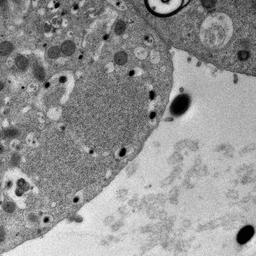

### i_143.jpg

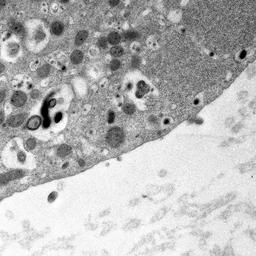
