## Supplementary figures and images for "Novel Automatic Classification of Human Adult Lung Alveolar Type II Cells Infected with SARS-CoV-2 through Deep Transfer Learning Approach"

### c_1.jpg

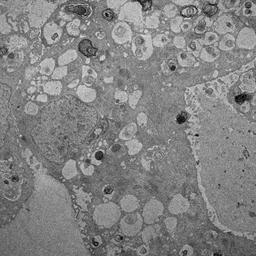

### c_2.jpg

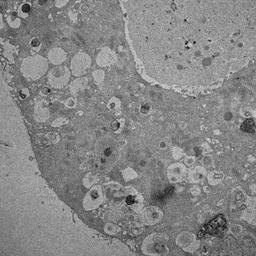

### c_3.jpg

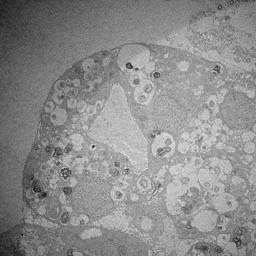

### c_4.jpg

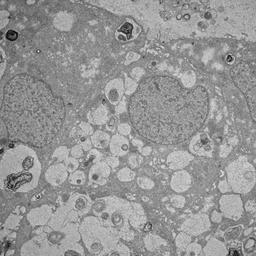

### c_5.jpg

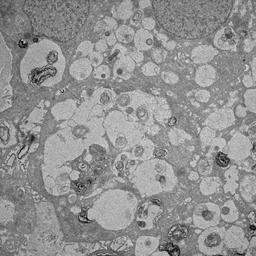

### c_6.jpg

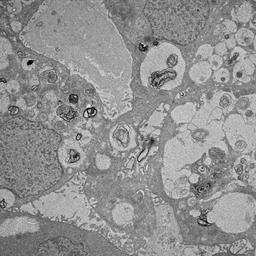

### c_10.jpg

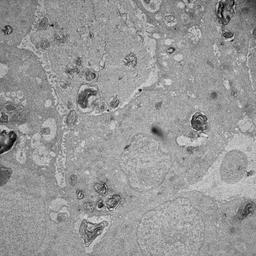

### c_11.jpg

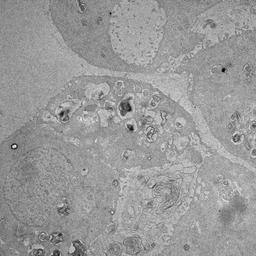

### c_12.jpg

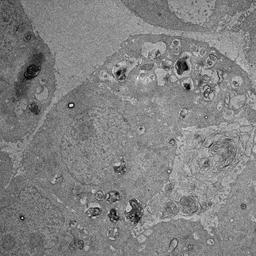

### c_13.jpg

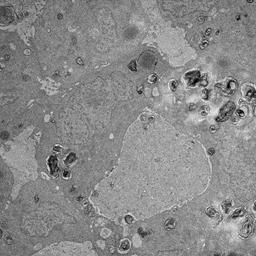

### c_14.jpg

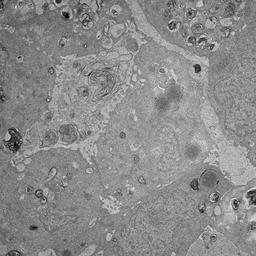

### c_15.jpg

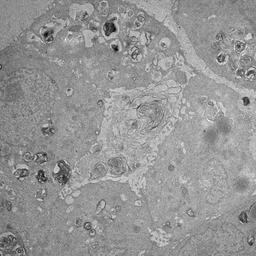

### c_16.jpg

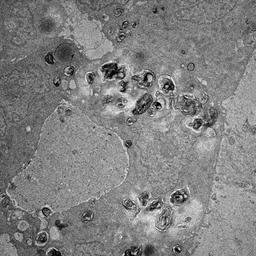

### c_17.jpg

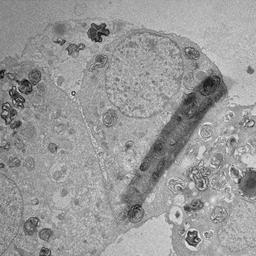

### c_18.jpg

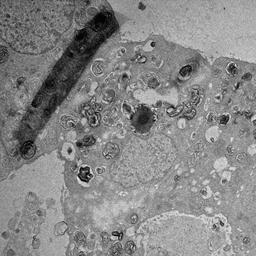

### c_19.jpg

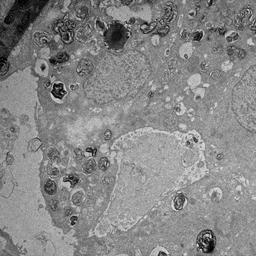

### c_20.jpg

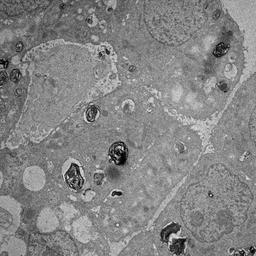

### c_21.jpg

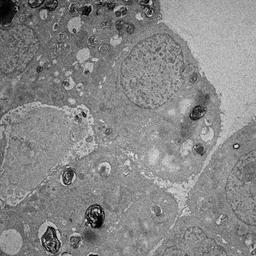

### c_22.jpg

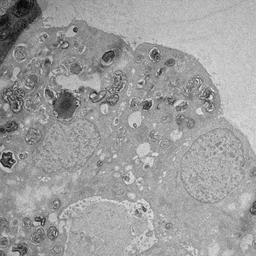

### c_23.jpg

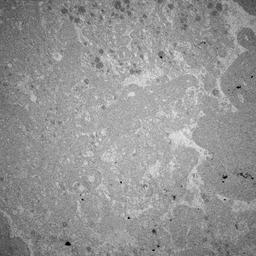

### c_24.jpg

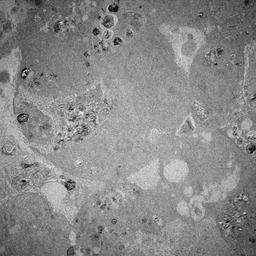

### c_25.jpg

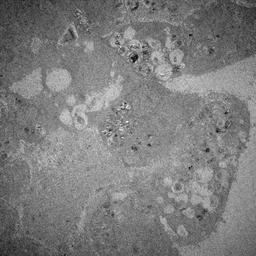

### c_26.jpg

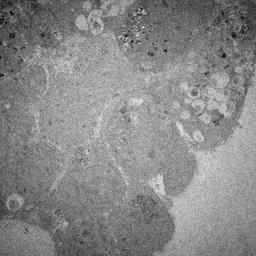

### c_27.jpg

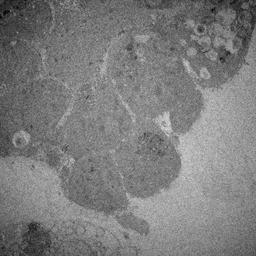

### c_28.jpg

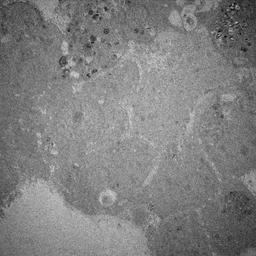

### c_29.jpg

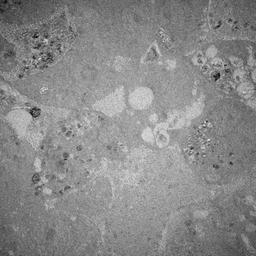

### c_30.jpg

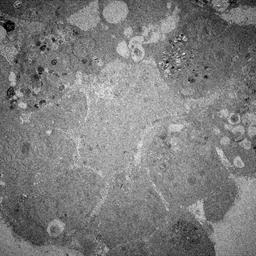

### c_31.jpg

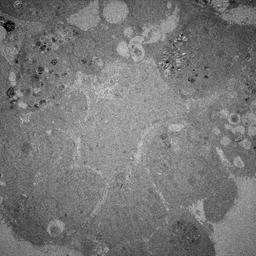

### c_32.jpg

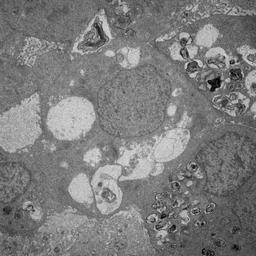

### c_33.jpg

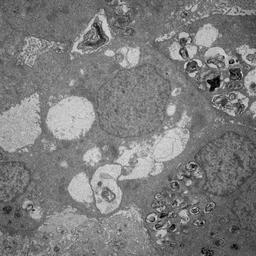

### c_34.jpg

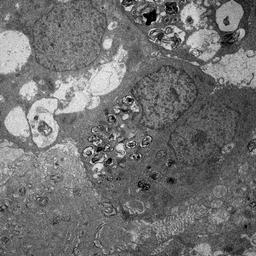

### c_35.jpg

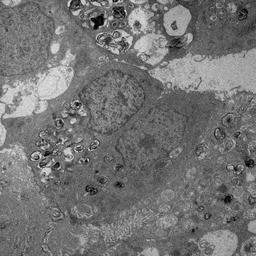

### c_36.jpg

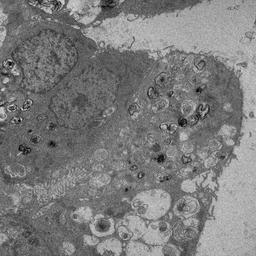

### c_37.jpg

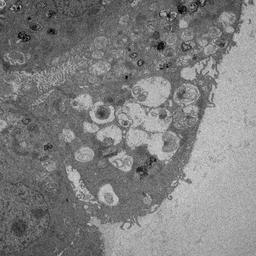

### c_38.jpg

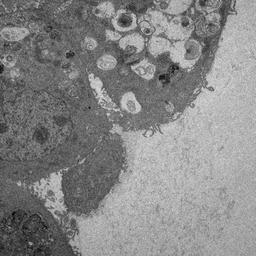

### c_39.jpg

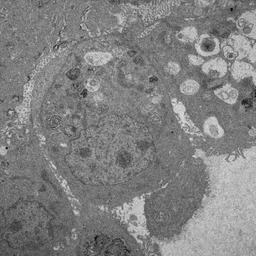

### c_40.jpg

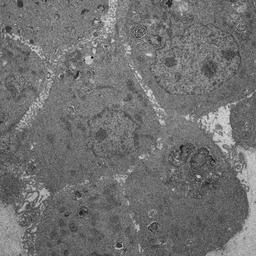

### c_41.jpg

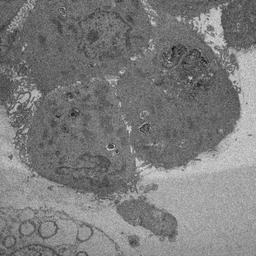

### c_42.jpg

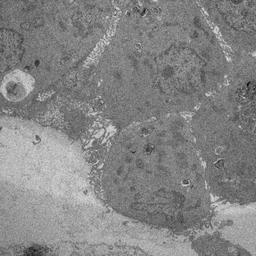

### c_43.jpg

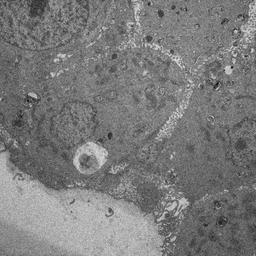

### c_44.jpg

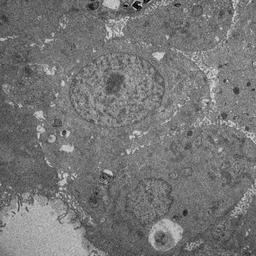

### c_45.jpg

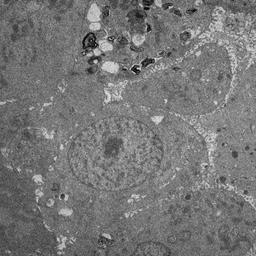

### c_46.jpg

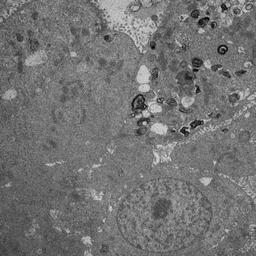

### c_47.jpg

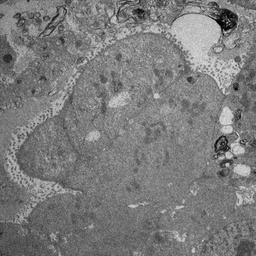

### c_48.jpg

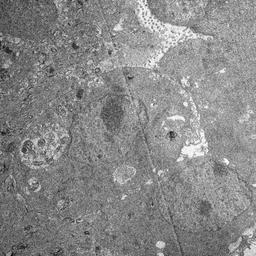

### c_49.jpg

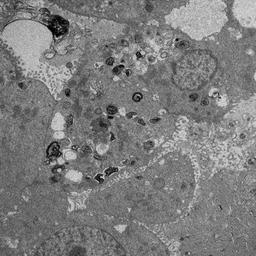

### c_50.jpg

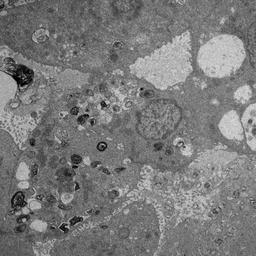

### c_51.jpg

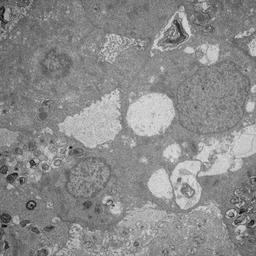

### c_52.jpg

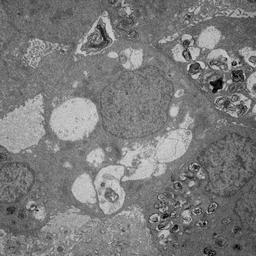

### c_53.jpg

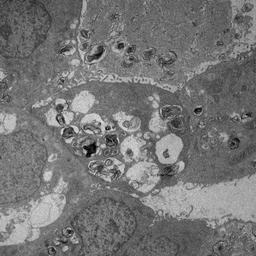

### c_54.jpg

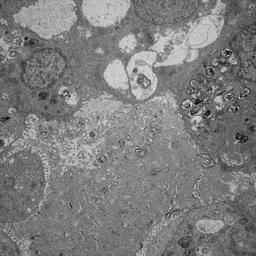

### c_55.jpg

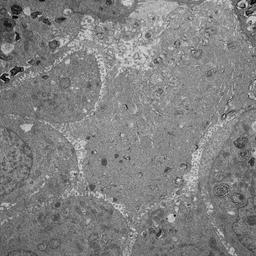

### c_56.jpg

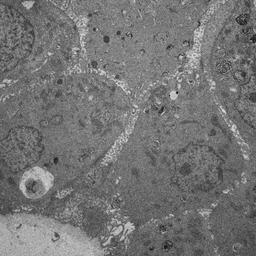

### c_57.jpg

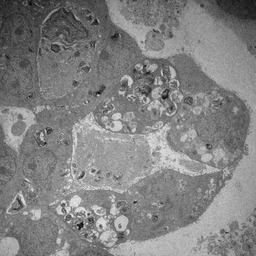

### c_58.jpg

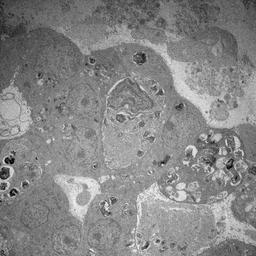

### c_59.jpg

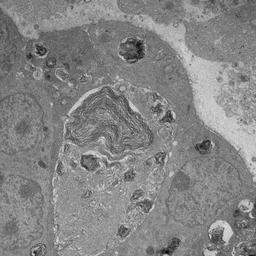

### c_100.jpg

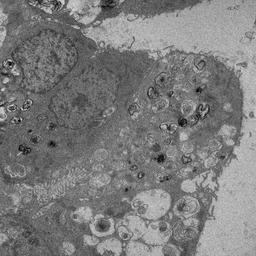

### c_101.jpg

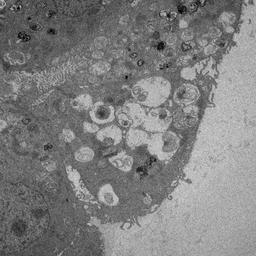

### c_102.jpg

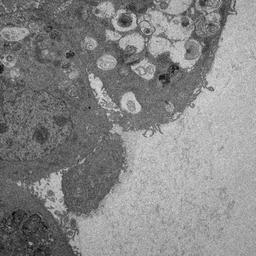

### c_103.jpg

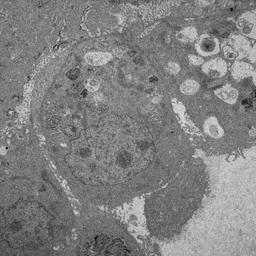

### c_104.jpg

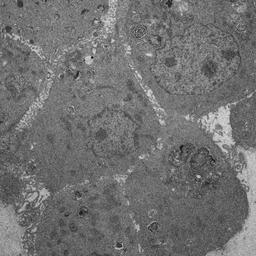

### c_105.jpg

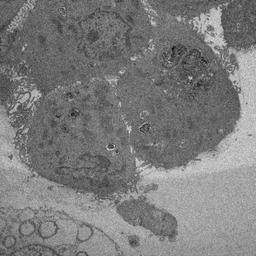

### c_106.jpg

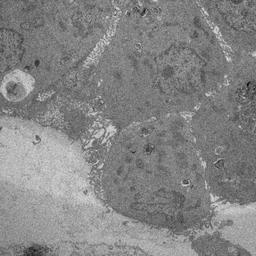

### c_107.jpg

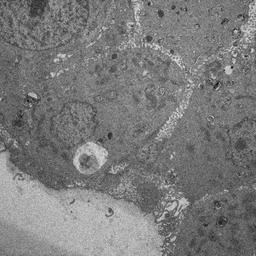

### c_108.jpg

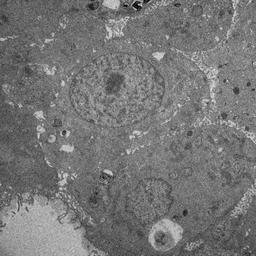

### c_109.jpg

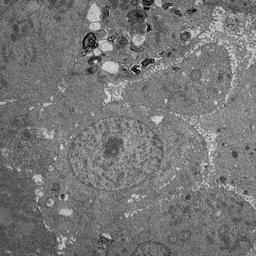

### c_110.jpg

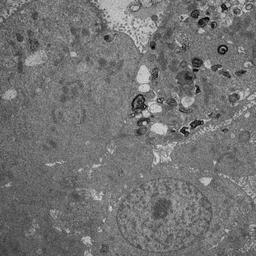

### c_111.jpg

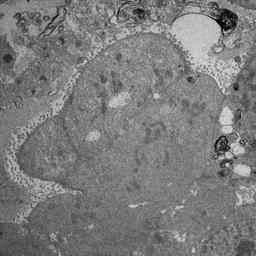

### c_112.jpg

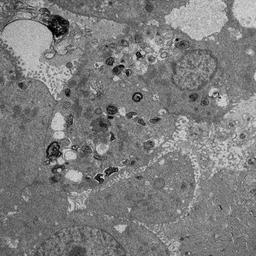

### c_113.jpg

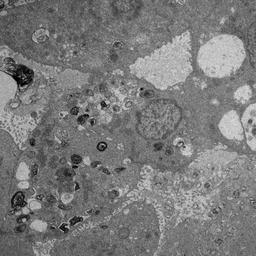

### c_114.jpg

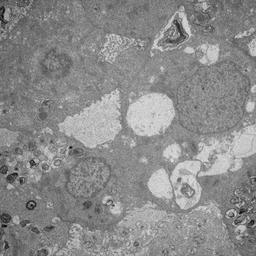

### c_115.jpg

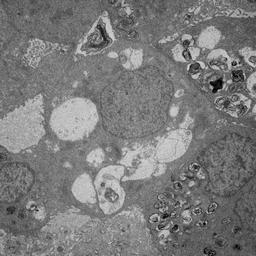

### c_116.jpg

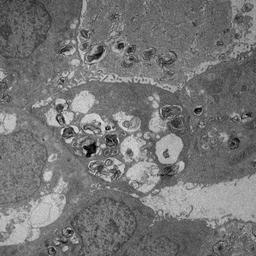

### c_117.jpg

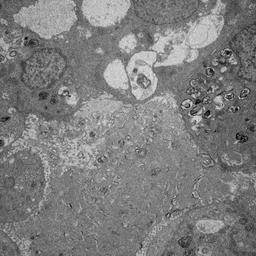

### c_118.jpg

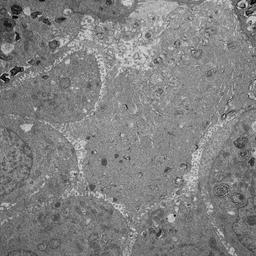

### c_119.jpg

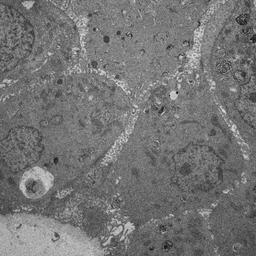

### c_120.jpg

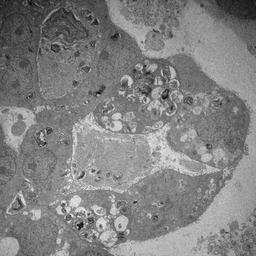

### c_121.jpg

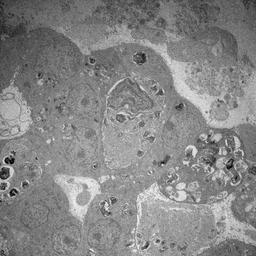

### c_122.jpg

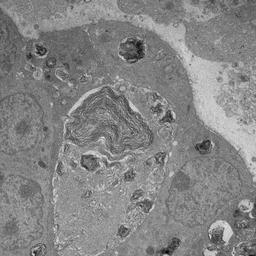

### c_123.jpg

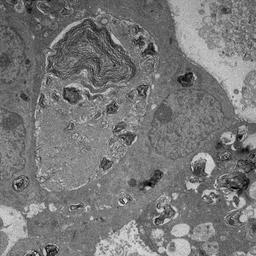

### c_124.jpg

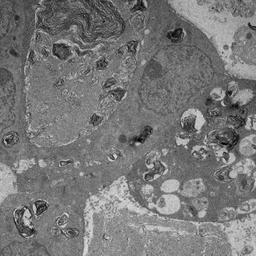

### c_125.jpg

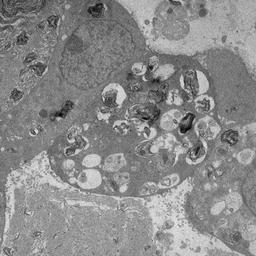

### c_126.jpg

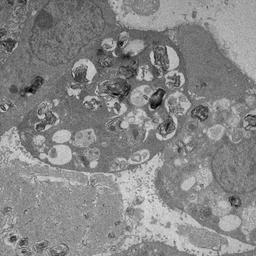

### c_127.jpg

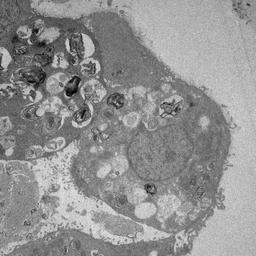

### c_128.jpg

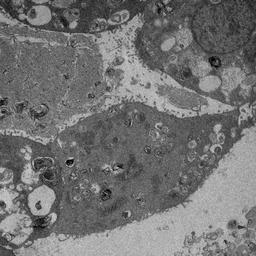

### c_129.jpg

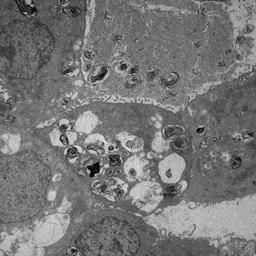

### c_130.jpg

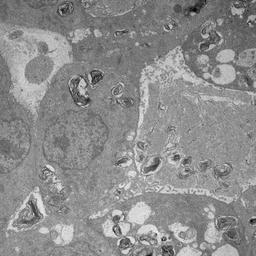

### c_131.jpg

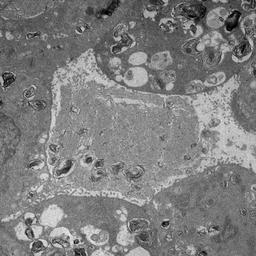

### c_132.jpg

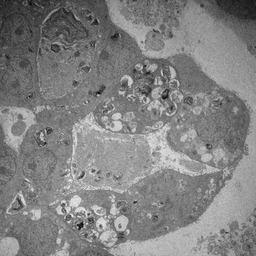

### c_133.jpg

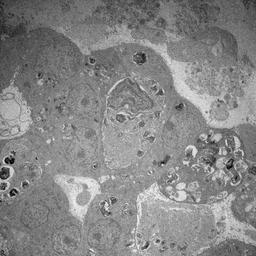

### c_134.jpg

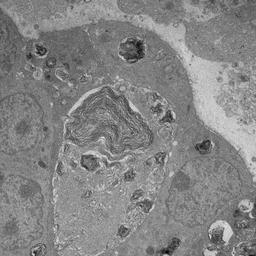

### c_135.jpg

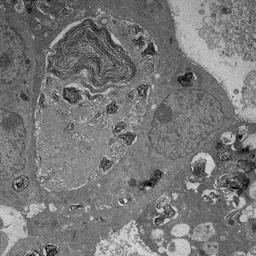

### c_136.jpg

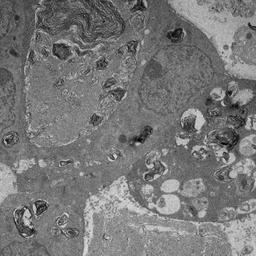

### c_137.jpg

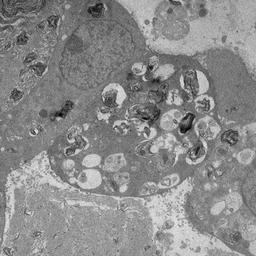

### c_138.jpg

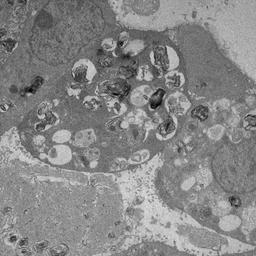

### c_139.jpg

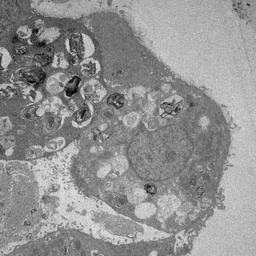

### c_140.jpg

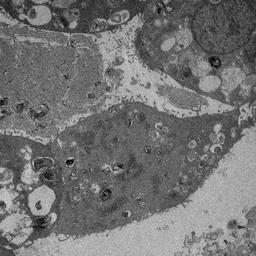

### c_141.jpg

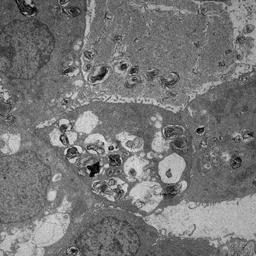

### c_142.jpg

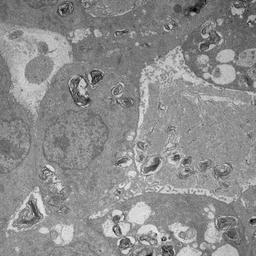

### c_143.jpg

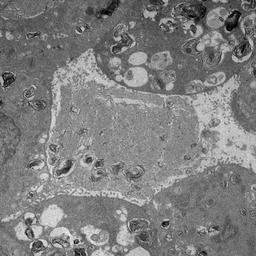
